# Forelimb movements contribute to hindlimb cutaneous reflexes during locomotion in cats

**DOI:** 10.1101/2024.03.13.584812

**Authors:** Jonathan Harnie, Rasha Al Arab, Stephen Mari, Sirine Yassine, Oussama Eddaoui, Pierre Jéhannin, Johannie Audet, Charly Lecomte, Christian Iorio-Morin, Boris I. Prilutsky, Ilya A. Rybak, Alain Frigon

## Abstract

During quadrupedal locomotion, central circuits interacting with somatosensory feedback coordinate forelimb and hindlimb movements. How this is achieved is not clear. To determine if forelimb movements modulate hindlimb cutaneous reflexes involved in responding to an external perturbation, we stimulated the superficial peroneal nerve in six intact cats during quadrupedal locomotion and during hindlimb-only locomotion (with forelimbs standing on stationary platform) and in two spinal-transected cats during hindlimb-only locomotion. We compared cutaneous reflexes evoked in six ipsilateral and four contralateral hindlimb muscles. Results showed similar occurrence and phase-dependent modulation of short-latency inhibitory and excitatory responses during quadrupedal and hindlimb-only locomotion in intact cats. However, the depth of modulation was reduced in the ipsilateral semitendinosus during hindlimb-only locomotion. Additionally, longer-latency responses occurred less frequently in extensor muscles bilaterally during hindlimb-only locomotion while short-latency inhibitory and longer-latency excitatory responses occurred more frequently in the ipsilateral and contralateral sartorius anterior, respectively. After spinal transection, short-latency inhibitory and excitatory responses were similar to both intact conditions, while mid- or longer-excitatory responses were reduced or abolished. Our results suggest that the absence of forelimb movements suppresses inputs from supraspinal structures and/or cervical cord that normally contribute to longer-latency reflex responses in hindlimb extensor muscles.

**NEW & NOTEWORTHY:** During quadrupedal locomotion, the coordination of forelimb and hindlimb movements involves central circuits and somatosensory feedback. To demonstrate how forelimb movement affects hindlimb cutaneous reflexes during locomotion, we stimulated the superficial peroneal nerve in intact cats during quadrupedal and hindlimb-only locomotion, as well as in spinal-transected cats during hindlimb-only locomotion. We show that forelimb movement influences the modulation of hindlimb cutaneous reflexes, particularly the occurrence of long-latency reflex responses.

## INTRODUCTION

Proper integration of cutaneous feedback within central neural circuits is required to execute specific motor tasks and perform necessary movement corrections. For instance, during locomotion, when the foot dorsum contacts an obstacle during the swing phase, a coordinated reflex response, termed the stumbling corrective reaction, allows the perturbed limb to step away from and over the obstacle to avoid falling, as shown in cats and humans (Forssberg et al. 1977; Forssberg 1979; Frigon et al. 2021; Haridas and Zehr 2003; Lecomte et al. 2023; Merlet et al. 2022; Prochazka et al. 1978; Quevedo et al. 2005a; Schillings et al. 1996; Van Wezel et al. 1997; Zehr et al. 1997). Electrically stimulating cutaneous nerves of the foot also elicits a coordinated reflex response, consistent with the stumbling corrective reaction in cats and humans (Buford and Smith 1993; Forssberg 1979; Haridas and Zehr 2003; Lam et al. 2003; Potocanac et al. 2016; Prochazka et al. 1978; Quevedo et al. 2005a, 2005b; Schillings et al. 2000, 2005; Van Wezel et al. 1997; Wand et al. 1980; Zehr et al. 1997). In cats, during the swing phase, short- (P1; latency of ∼8-12 ms) and mid- (P2; latency of ∼20-25 ms) latency excitatory responses in ipsilateral hindlimb (i.e. the limb stimulated) muscles that flex the knee and ankle joints are observed following electrical stimulation, resulting in limb flexion. To maintain balance, the activity of extensor muscles of the contralateral hindlimb (i.e. the limb opposite stimulation) is reinforced via mid-latency (P2) excitatory responses. The same stimulus applied during stance generates a short-latency inhibitory response (N1; latency of ∼8-12 ms) in ipsilateral extensor muscles to briefly pause stance, referred to as the stumbling preventive reaction, that are followed by mid- (P2) or long-latency (P3; latency of ∼35 ms) responses (Merlet et al. 2022; Quevedo et al. 2005b). Phase-dependent modulation ensures that cutaneous reflexes are functionally relevant (Zehr and Stein 1999). Reflex responses evoked by stimulating nerves of the foot are also observed in arm/forelimb muscles in cats and humans (Haridas and Zehr 2003; Hurteau et al. 2018; Mari et al. 2023; Miller et al. 1977; Zehr et al. 2001). Thus, cutaneous reflexes of each foot are distributed to the four limbs and are part of a whole body response to a perturbation (Merlet et al. 2022).

During quadrupedal locomotion in terrestrial mammals and bipedal locomotion in humans, all four limbs contribute to dynamic stability and proper coordination. A bi-directional influence between the arms and legs has been demonstrated in humans by showing that the position or rhythmic movement of the arms or legs can influence the activity of the legs or arms, respectively (Delwaide et al. 1973, 1977; Eke-Okoro 1994; Ferris et al. 2006; Frigon et al. 2004; Haridas and Zehr 2003; Hiraoka 2001; Hiraoka and Nagata 1999; Hundza and Zehr 2009; Kagamihara et al. 2003; Kao and Ferris 2005; Meinck and Piesiur-Strehlow 1981; Sakamoto et al. 2006, 2007; Zehr and Duysens 2004). For instance, in humans, stretch or tendon tap reflexes in leg muscles are modified by the position of the arm (Delwaide et al. 1977). Rhythmic arm cycling reduced soleus H-reflex amplitude in humans when the legs are in a static position (Frigon et al. 2004). Passive shoulder flexion-extension movements facilitated the soleus H-reflex (Hiraoka and Nagata 1999), whereas active arm swinging movements reduced its amplitude (Hiraoka 2001). Using a transverse split-belt treadmill (unequal speed of the fore- and hindlimbs), studies found that movement of the forelimbs influenced the activity of the hindlimbs and vice-versa in decerebrate cats (Akay et al. 2006), as well as in intact adult cats and kittens (Cruse and Warnecke 1992; Thibaudier et al. 2013; Thibaudier and Frigon 2014). Interestingly, these bidirectional influences appear asymmetrical, with a greater influence of the forelimb/arms on the hindlimb/legs in cats (Akay et al. 2006; Hurteau et al. 2018; Thibaudier et al. 2013) and humans (Sakamoto et al. 2014). How forelimb movements contribute to reflex modulation in the hindlimbs of quadrupeds is not known.

We recently compared the hindlimb pattern of intact cats during quadrupedal treadmill locomotion and during hindlimb-only locomotion where the forelimbs were placed on a stationary platform as well as cats following a spinal transection (i.e. spinal cats) during hindlimb-only locomotion (Harnie et al. 2022). We found that the spatiotemporal pattern as well as the activity of hindlimb muscles during hindlimb-only locomotion in spinal cats more closely resembled the pattern during hindlimb-only locomotion of intact cats. Thus, in intact cats, forelimb movements shape the hindlimb locomotor pattern. The goal of the present study was to investigate the contribution of forelimb movements on hindlimb cutaneous reflexes in intact cats by comparing responses during quadrupedal locomotion and hindlimb-only locomotion. In the second part of this study, we compared these data with hindlimb cutaneous reflexes obtained after a complete spinal transection. We hypothesized that forelimb movements contribute to the modulation of hindlimb cutaneous reflexes during locomotion.

## MATERIALS AND METHODS

### Animals and ethical information

All procedures were approved by the Animal Care Committee of the Université de Sherbrooke and were in accordance with policies and directives of the Canadian Council on Animal Care (Protocol 442-18). Six adult cats, 4 males and 2 females, weighing between 3.7 kg and 6.5 kg (5.0 ± 1.0) were used in the present study. We followed ARRIVE guidelines for animal studies (Percie du Sert et al. 2020). Before and after experiments, cats were housed and fed in a dedicated room within the animal care facility of the Faculty of Medicine and Health Sciences at the Université de Sherbrooke. As part of our effort to maximize the output of each animal, cats were used in other studies to answer different scientific questions, many of which are in preparation.

### Surgical procedures, electrode implantation and spinal transection

Surgeries were performed under aseptic conditions with sterilized instruments in an operating room. Before surgery, cats first received an intramuscular injection containing ketamine (0.05 ml/kg), Butorphanol (0.4 ml/kg), Dexmedetomidine (0.06 ml/kg). Once sedation was visible, the cat was placed on a heating mat and covered with a blanket. A long-lasting Optix-care eye gel (hyaluronic acid) was administered. The bladder was emptied by manual pressure and vital signs were monitored, including heart rate, respiratory rate, mucous membrane color and body temperature, as well as depth of sedation. In the surgical room, cats were then anesthetized with isoflurane (1.5-3%) and O_2_ (1 L/min) using a mask and then intubated with a flexible endotracheal tube. Cats received a continuous infusion of lactated Ringers solution (3 ml/kg/h) during the surgery through a catheter placed in a cephalic vein. Anesthesia was maintained by adjusting isoflurane concentration as needed and by monitoring cardiac and respiratory rates. Body temperature was monitored with a rectal thermometer and maintained within physiological range (38 ± 0.5°C) using a water-filled heating pad placed under the animal and an infrared lamp ∼50 cm over it. The depth of anesthesia was confirmed by applying pressure to a paw (to detect limb withdrawal) and by assessing the size and reactivity of pupils. Before surgery, the animal’s skin was cleaned with chlorhexidine soap and meloxicam (0.2 mg/kg), an anti-inflammatory agent, was injected subcutaneously along with an anesthetic block at the incision sites (2:1 Bupivacaine 0.25% / Lidocaine 2.0%). After surgery, we taped a transdermal fentanyl patch (25 µg/h) to the back of the animal 2-3 cm rostral to the base of the tail for prolonged analgesia (removed after 5-7 days). Cats were placed in an incubator until they regained consciousness. Meloxicam and Chloramphenicol (Chlor-palm), an anti-inflammatory and an antibiotic, were given orally for 4 and 5 consecutive days, respectively. At the conclusion of the experiments, cats received a lethal dose (120 mg/kg) of sodium pentobarbital through the cephalic vein.

#### Electrode implantation

To record the electrical activity of muscles (EMG, electromyography), a skin incision was made at the midline of the skull and muscles and fascia were retracted laterally. Pairs of Teflon-insulated multistrain fine wires (AS633; Cooner Wire, Chatsworth, CA) were directed subcutaneously from two head-mounted 34-pin connectors (Omnetics Connector, Minneapolis, MN). Skin and fascial incisions were made to expose selected hindlimb muscles and electrodes were sewn into the belly for bipolar recordings, with 1-2 mm of insulation stripped from each wire. The head connector was secured to the skull using six screws and dental acrylic. Electrode placement was verified during surgery by electrically stimulating each muscle through the appropriate head connector channel. The current data set includes EMG from the following muscles bilaterally: soleus (SOL, ankle extensor), vastus lateralis (VL, knee extensor), biceps femoris anterior (BFA, hip extensor), anterior sartorius (SRT, hip flexor/knee extensor), semitendinosus (ST, knee flexor/hip extensor) and tibialis anterior (TA, ankle flexor). For nerve stimulation, pairs of Teflon-insulated multistrain fine wires (AS633; Cooner Wire) were passed through a silicon tubing. A horizontal slit was made in the tubing and wires within the tubing were stripped of their insulation. The ends protruding through the cuff were knotted to hold the wires in place and glued. The ends of the wires away from the cuff were inserted into four-pin connectors (Hirose or Samtec) for bipolar nerve stimulation. Cuff electrodes were directed subcutaneously from head-mounted connectors and placed around the left and right SP nerves at ankle level. At this level, the SP nerve is purely cutaneous (Bernard et al. 2007).

#### Spinal transection

The skin was incised over the last thoracic vertebrae and after carefully setting aside muscle and connective tissue, a small dorsal laminectomy was made. After exposing the spinal cord, xylocaine (lidocaine hydrochloride, 2%) was applied topically followed by two to three intraspinal injections. The spinal cord was then completely transected with surgical scissors between the 12th and 13th thoracic vertebrae. We cleaned the 0.5-cm gap between the two cut ends of the spinal cord and stopped any residual bleeding. We verified that no spinal cord tissue remained connecting rostral and caudal ends, which we later confirmed histologically. A hemostatic agent (Spongostan) was placed within the gap, and muscles and skin were sewn back to close the opening in anatomic layers. After spinal transection, we manually expressed the cat’s bladder and large intestine one to two times daily, or as needed.

### Data collection and analysis

The main goal was to assess the modulation of cutaneous hindlimb reflexes during quadrupedal and hindlimb-only locomotion in the intact state. In two cats, we also assessed the modulation of hindlimb cutaneous reflexes during hindlimb-only locomotion six weeks after spinal transection. During experiments, cats performed tied-belt (equal left-right speeds) locomotion in quadrupedal and hindlimb-only conditions on a treadmill with two independently controlled running surfaces 120 cm long and 30 cm wide (Bertec) at a speed of 0.4 m/s. In the hindlimb-only condition, the forelimbs were placed on a stationary platform. In the spinal state, we collected data with perineal stimulation. For perineal stimulation, an experimenter manually pinched the skin under the tail with the index finger and thumb. We did not provide weight support, although an experimenter gently held the tail to provide equilibrium. Videos of the left and right sides were obtained with two cameras (Basler AcA640-100g) at 60 frames/s with a spatial resolution of 640 by 480 pixels. A custom-made program (Labview) acquired the images and synchronized them with EMG data. EMG signals were pre-amplified (10×, custom-made system), band-pass filtered (30-1000 Hz) and amplified (100-5000×) using a 16-channel amplifier (AM Systems Model 3500). EMG data were digitized (5000 Hz) with a National Instruments card (NI 6032E), acquired with custom-made acquisition software, and stored on a computer. We electrically stimulated the SP nerve with a Grass S88 Stimulator at an intensity of 1.2 times the motor threshold, defined as the voltage required to elicit a small consistent short-latency (∼10 ms) excitatory EMG response in an ipsilateral flexor, such as semitendinosus (ST). We determined the motor threshold during quadrupedal locomotion and applied the same stimulation intensity during hindlimb-only locomotion. This stimulation intensity mainly activates large diameter Aβ afferents (Drew and Rossignol 1987; LaBella et al. 1992; Pratt et al. 1991; Quevedo et al. 2005a, 2005b). Each stimulation consisted of a train of three 0.2 ms pulse duration at 300 Hz frequency. During a trial, ∼120 stimuli were delivered pseudo-randomly every two to four locomotor cycles at varying delays relative to an ipsilateral hindlimb extensor burst onset to evoke responses at different times during the step cycle. We obtained a minimum of five stimuli per bin in each condition for all cats. Each locomotor condition tested lasted 6 to 9 min. For a given nerve stimulation, we collected data for both locomotor conditions within a single session and at the same stimulation intensity.

Kinematic data were analyzed as described previously (Harnie et al. 2018, 2021, 2022). By visual detection, we determined hindlimb contact as the first frame where the paw made visible contact with the treadmill surface, and limb liftoff as the most caudal displacement of the toe. Cycle duration corresponded to the interval of time between consecutive paw contacts of the same limb. Stance duration was defined as the interval of time from initial contact to liftoff of the same limb. Swing duration was calculated as the difference between cycle duration and stance duration. We measured the proportion of the stance and swing phases as a percentage of cycle duration. We measured the temporal and spatial symmetry indexes because left-right symmetry has been shown to influence reflex modulation (Hurteau et al. 2017; Mari et al. 2023). We measured the temporal phase interval by dividing the interval of time between stance onsets of the right and left hindlimbs by right hindlimb cycle duration. For the gap interval, we first measured step length as the distance between the leading and trailing limb at stance onset of the leading limb and stride length as the horizontal distance traveled from stance offset to onset plus the distance traveled by the contralateral treadmill belt during the swing phase. Then, we divided right hindlimb step length by right hindlimb stride length. For the temporal symmetry index, we measured the absolute deviation of the temporal phase interval from a perfect symmetry of 0.5. For the spatial symmetry index, we measured the absolute deviation of the gap interval from a perfect symmetry of 0.5. Symmetry indexes were then multiplied by 100 and expressed as a percentage.

Reflex data were analyzed as described previously (Hurteau et al. 2017, 2018; Hurteau and Frigon 2018). We removed sections from analysis where the cat stepped irregularly or removed its forelimbs from the platform in the hindlimb-only condition based on EMG and video data. The step cycle was synchronized to the soleus extensor burst onset (SOL) in the stimulated limb (the ipsilateral limb). Cycles were tagged as stimulated (i.e., cycles with stimulation) or control (i.e., cycles without stimulation) and divided into 10 subphases (bins) of equal duration. Control cycles were averaged, rectified and separated into 10 subphases to provide a baseline locomotor EMG (blEMG) in each bin. Stimulated cycles were averaged, rectified and separated in each bin according to the time when stimulation occurred within the cycle. To quantify reflex responses, a window of 80 ms around the stimulation was opened and the blEMG was superimposed on the EMG averaged from stimulated cycles for the 10 bins. The same experimenter (Harnie) determined onsets and offsets responses by visual inspection, defined as prominent positive or negative deflections away from the blEMG, using previous studies as guidelines (Duysens and Loeb 1980; Duysens and Stein 1978; Hurteau et al. 2017, 2018; Hurteau and Frigon 2018; Loeb 1993; Pratt et al. 1991) and 97.5% confidence intervals. Short-latency excitatory (P1) and inhibitory (N1) responses in ipsilateral and contralateral limbs started at 9-18 ms following stimulation. Mid-latency (20–25 ms) excitatory and inhibitory responses are termed P2 and N2 responses, respectively. Long-latency (35 ms) excitatory and inhibitory responses are termed P3 and N3 responses, respectively. We found N2 and N3 responses inconsistently and excluded them from further analysis.

The EMG of reflex responses were integrated and then subtracted from the integrated blEMG in the same time window to provide a net reflex value. This net reflex value was then divided by the integrated blEMG value to determine if reflex modulation is independent from blEMG activity (Frigon et al. 2009; Frigon and Rossignol 2007, 2008a; Hurteau et al. 2017, 2018; Hurteau and Frigon 2018; Matthews 1986). Phase and condition (quadrupedal and hindlimb-only locomotion) modulation was illustrated by normalizing reflex responses in each muscle to the maximal value (expressed as a percentage) obtained in one of the two locomotor conditions. To determine the effect of the locomotor condition on the phase-dependent modulation, we obtained a reflex index by measuring the difference between the largest and smallest responses out of the 10 bins for each locomotor condition (Hurteau et al. 2017, 2018; Hurteau and Frigon 2018).

To characterize EMG bursts, we selected 15–30 control cycles. The onset and offset of bursts in the selected muscles were identified through visual examination (Harnie) using a custom-made program applied to the raw EMG waveforms. The burst durations were then expressed as a percentage of the cycle duration. Mean EMG amplitude was measured by integrating the full-wave rectified EMG burst from onset to offset and dividing it by its burst duration. Subsequently, we normalized this value by the maximal value obtained in one of the two locomotor conditions.

### Statistical analysis

We quantified reflex responses in the ipsilateral and contralateral SOL, VL, BFA, SRT, ST and TA. In five of six intact cats, both the left and right SP nerves evoked responses. We treated responses evoked by left and right nerves separately and pooled them for statistical analysis (n = 11 nerves stimulated in six cats). To compare the occurrence of responses in the two locomotor conditions, we performed a chi-square test on the presence or not of reflex responses across bins. To assess phase-dependent modulation, we performed a nonparametric one-factor (bins) Friedman test on short-, mid- and longer-latency responses (N1, P1, P2/P3) for pooled data. In muscles that showed P2 and P3 responses, we pooled them because we believe they are mediated by the same pathway and depend on N1 duration. We used the Wilcoxon signed-rank test on pooled data to assess the effect of locomotor condition on the proportion of stance and swing durations, EMG burst durations and amplitudes, as well as symmetry indexes and reflex modulation indexes. Statistical significance for all tests was set at P < 0.05.

## RESULTS

### Modulation of the hindlimb pattern during quadrupedal and hindlimb-only locomotion

In the current study, we investigated cutaneous reflexes during quadrupedal and hindlimb-only locomotion in intact cats at a treadmill speed of 0.4 m/s. We recently published a study characterizing the locomotor pattern, including electromyography (EMG) in selected hindlimb muscles, in the two locomotor conditions over a range of treadmill speeds (Harnie et al. 2022), which revealed significant differences in spatiotemporal parameters, such as reduced cycle and stance durations, as well as shortened step and stride lengths during hindlimb-only locomotion compared to quadrupedal locomotion. During hindlimb-only locomotion, we also observed changes in EMG activity, such as a longer semitendinosus (ST) burst and double bursts in the iliopsoas and anterior sartorius (SRT). We confirm these changes in EMG and cycle/stance durations in the present study, as shown for a representative cat (**Fig. 1A**). Note the shorter cycle and stance durations as well as double bursts in the SRT and tibialis anterior (TA) during hindlimb-only locomotion. However, the proportions of stance (P = 0.249) and swing (P = 0.249) durations as a function of cycle duration were similar between hindlimb-only and quadrupedal locomotion (**Fig. 1B**). The normalized burst durations of vastus lateralis (VL) (P = 0.046; +7%) and biceps femoris anterior (BFA) (P = 0.046; +8%) were significantly longer during hindlimb-only locomotion, whereas the normalized durations of the swing-related burst of SRT (P = 0.028; -10%) and TA (P = 0.028; -7%) muscles were significantly shorter compared to quadrupedal locomotion (**Fig. 1C**). We found a significant decrease in EMG burst amplitude for soleus (SOL) (P = 0.028; - 10%) and SRT (P = 0.028; -34%) muscles during hindlimb-only locomotion compared to quadrupedal locomotion, while a significant increase was observed for VL (P = 0.028; +39%), ST (P = 0.046; +18%) and TA (P = 0.046; +20%), with no significant difference for BFA (P = 0.753) (**Fig. 1D**). We also measured temporal and spatial symmetry indexes during quadrupedal and hindlimb-only locomotion to assess the consistency of left-right coordination on a step-by-step basis between the two tasks and found no significant differences for temporal (P = 0.345) or spatial indexes (P = 0.917) (**Fig. 1E**).

**Figure 1.**
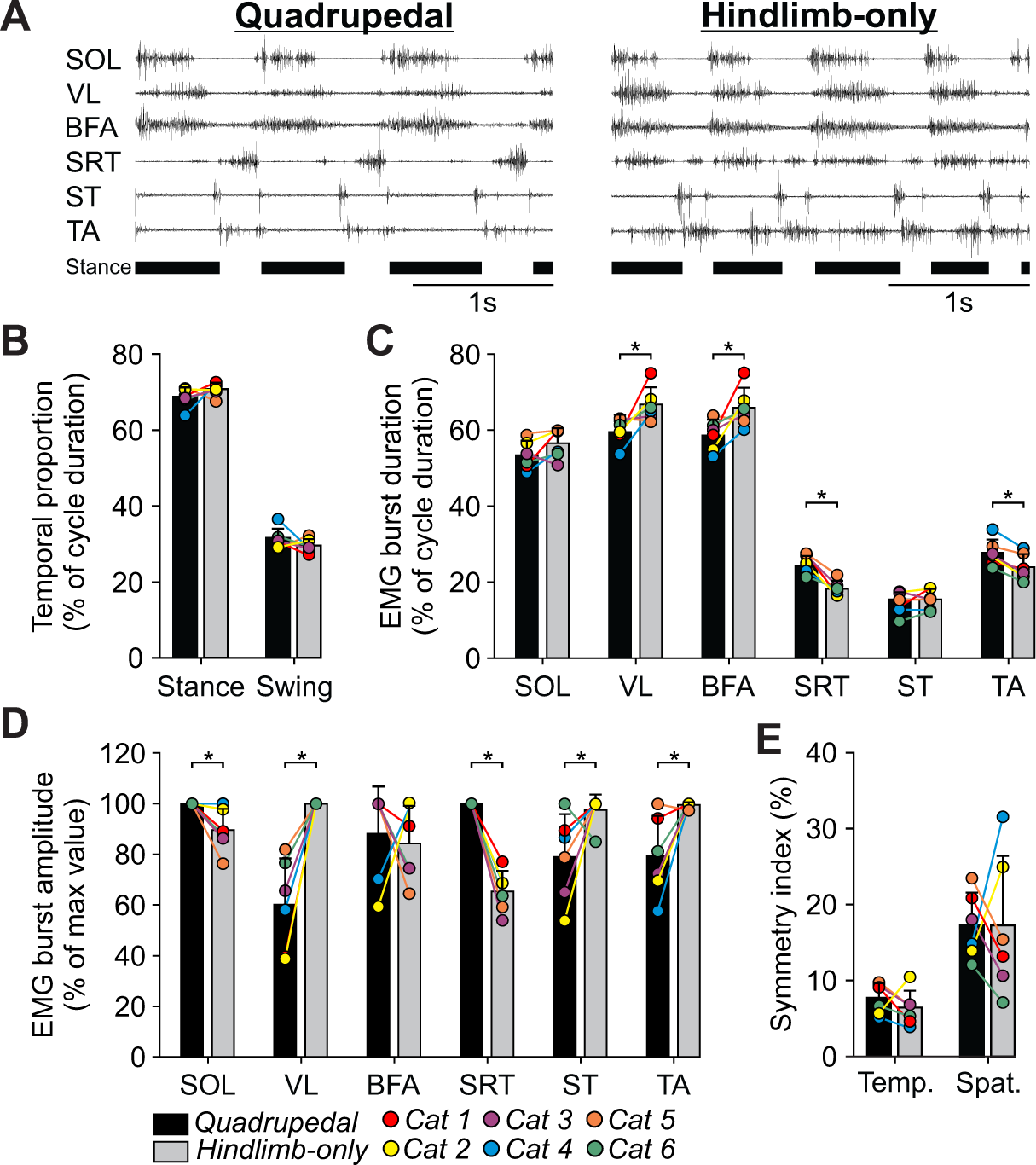
EMG activity and kinematic characteristics of quadrupedal and hindlimb-only locomotion in intact cats at 0.4 m/s. **A)** EMG activity from six hindlimb muscles along with stance phases (horizontal thick lines) of the right (R) hindlimb in one intact cat during quadrupedal and hindlimb-only locomotion. Data are from Cat 4. SOL, soleus; VL, vastus lateralis; BFA, biceps femoris anterior; SRT, anterior sartorius; ST, semitendinosus; TA, tibialis anterior. **B)** Stance and swing durations expressed as a percentage of cycle duration. **C)** EMG burst duration expressed as a percentage of cycle duration. Note that for SRT and TA, we only measured the flexor/swing-related burst. **D)** EMG burst amplitudes expressed as a percentage of the maximum value obtained in one of the two conditions. **E)** Temporal (Temp.) and spatial (Spat.) symmetry indexes, calculated by measuring the relative deviation of the phase and gap intervals from a perfect out-of-phase interval of 0.5, expressed as a percentage. Each bar is the mean ± standard deviation for the group (n = 6 cats). Individual data are shown with different colored circles. Asterisks represent significant effect of condition (Wilcoxon): *p<0.05.

### Cutaneous reflex responses in ipsilateral hindlimb muscles

**Table 1** shows the occurrence of ipsilateral and contralateral responses for each muscle with SP nerve stimulation. For a given muscle, the number of nerves stimulated (n = 11 in six cats) where N1, P1 or P2/P3 responses were observed is represented as a fraction of the total number of nerves stimulated, with the percentage indicated in parentheses. We also calculated the number of bins where responses were observed as a fraction of the total number of bins, with the percentage indicated in parentheses. We used Chi-square tests to determine if reflex response occurrence differed between locomotor conditions by comparing the number of bins with a response. To investigate the phase- and condition-dependent modulation of cutaneous reflexes, we stimulated the SP nerve at different times during the step cycle and recorded responses in ipsilateral and contralateral muscles in both tasks.

**Table 1.**
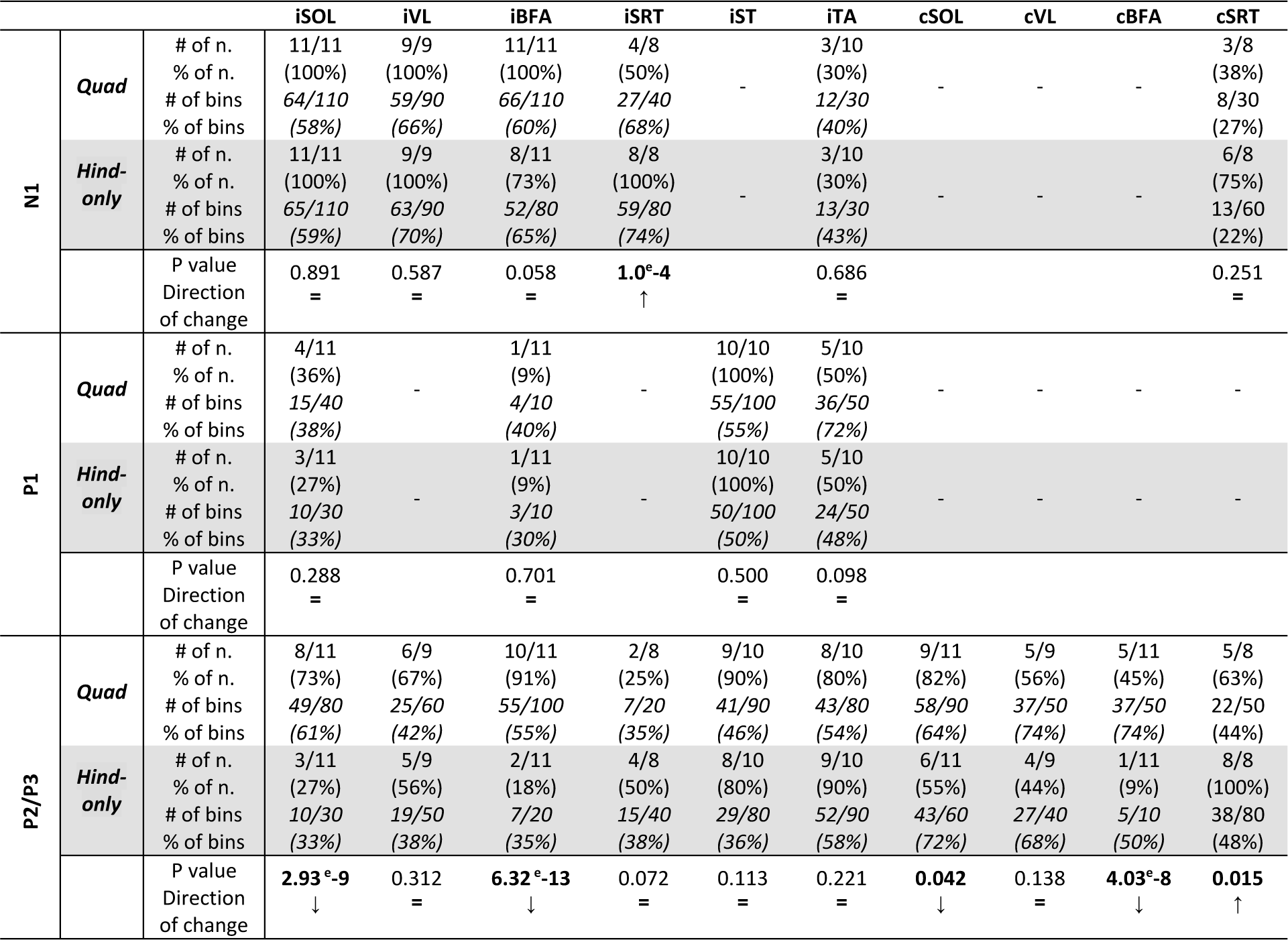
Occurrence of responses elicited by SP nerve stimulation in ipsilateral and contralateral hindlimb muscles during quadrupedal and hindlimb-only locomotion. For each muscle, the number of nerves stimulated (# of n.) where N1, P1 and P2/P3 responses were observed is represented as a fraction of the total number of nerve stimulated, with the percentage indicated in parentheses (% of n.). We also calculated the number of bins (# of bins) where responses were observed as a fraction of the total number of bins, with the percentage indicated in parentheses (% of bins). Chi-square test was performed between conditions on the number of bins where responses occurred. Up and down arrows indicate a significant increase or decrease, respectively in reflex response occurrence during hindlimb-only locomotion compared to quadrupedal locomotion. The equal sign represents no significant changes.

To facilitate comparisons between conditions and muscles, we organized **Figures 2-4** in the same way. In these figures, the two leftmost panels show reflex responses evoked in the 10 bins during quadrupedal and hindlimb-only locomotion for a single cat. To illustrate the deviation of the responses from the normal EMG pattern, the EMG from cycles that received stimulation (black traces) was averaged and superimposed on the averaged EMG from cycles without stimulation (the blEMG in grey). To the right of each panel, we show the average rectified EMG activity of the muscle from control cycles and the stance phase normalized to ipsilateral SOL onset. For each nerve stimulation, we normalized responses in each bin to the largest response obtained out of both conditions. Thus, responses in some bins are less visible because of this scaling but they were still analyzed. The scatter plots to the right of the individual cat data show normalized reflex response amplitudes for pooled data. Periods of muscle activity in both locomotor conditions are shown above the scatter plots in the two conditions.

**Figure 2.**
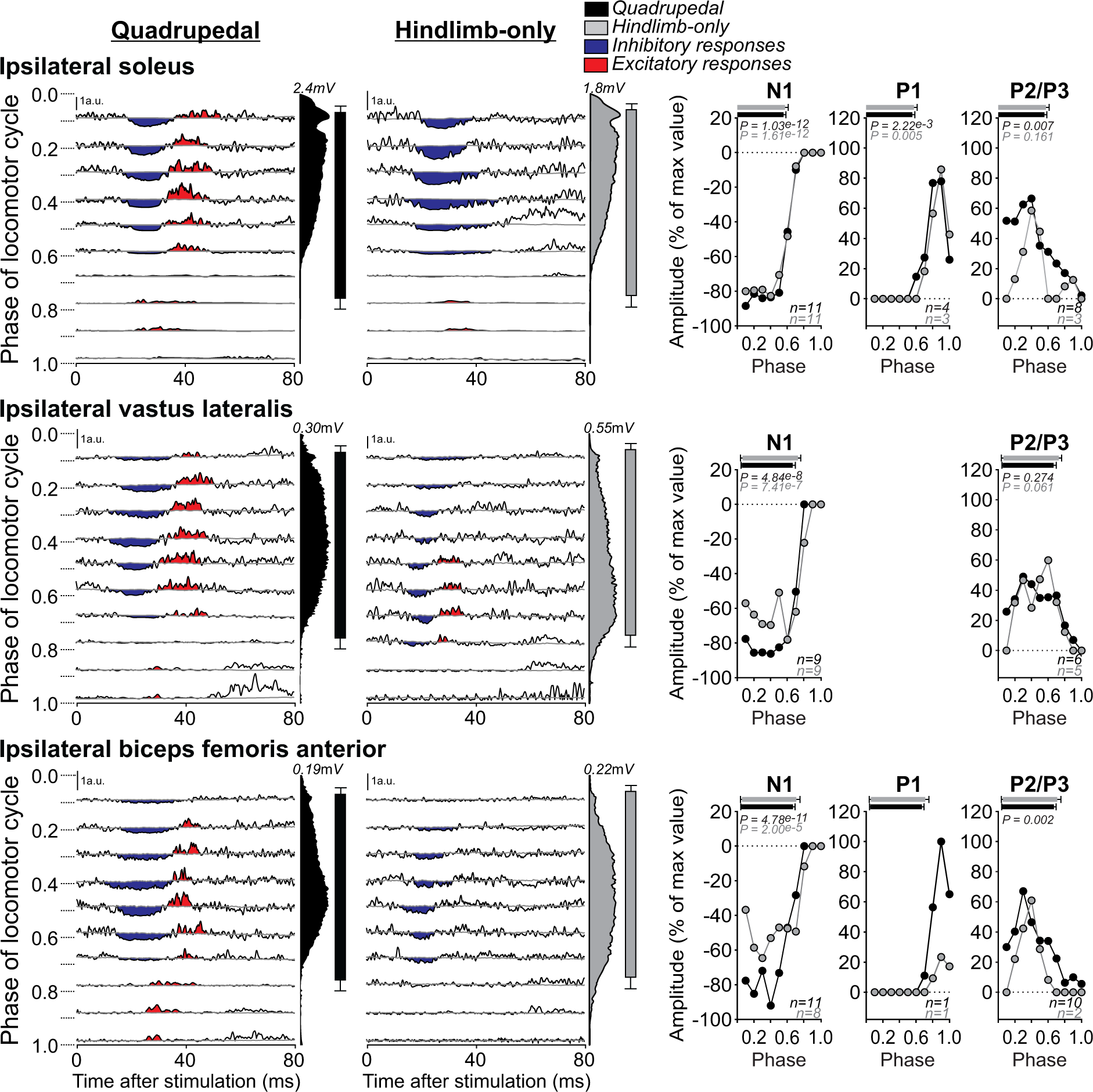
Phase- and condition-dependent modulation of cutaneous reflexes in ipsilateral extensor muscles. SP nerve stimulation evoked responses in the ipsilateral soleus (n = 11 nerves in 6 cats), vastus lateral (n = 9 nerves in 6 cats) and biceps femoris anterior (n = 11 nerves in 6 cats) during quadrupedal and hindlimb-only locomotion. Each panel represents averaged waveforms in the 10 bins of the cycle with a post-stimulation window of 80 ms in a single cat (Cat 3 for all three muscles). Black traces represent averaged cycles that received a stimulation (i.e., reflex responses, n = 7–18 stimuli per bin). Gray traces represent averaged cycles with no stimuli. Scale bars are shown in arbitrary units (a.u.) for both locomotor conditions. Blue and red areas represent inhibitory and excitatory responses, respectively. Aligned vertically is the average rectified EMG and the stance phase aligned to ipsilateral soleus muscle onset. Scatter plots represent reflex amplitudes (mean) in each bin of the cycle expressed as a percentage of the maximal response found in one of the bins for each nerve stimulated during quadrupedal (black circles) and hindlimb-only locomotion (gray circles). Short-latency (P1, N1) and mid/longer-latency (P2/P3) responses are shown separately. Horizontal bars at the top of the scatter plots represent the period of muscle activity for each locomotor condition (n = 7-15 control cycles) for pooled data. P values at the top of the scatter plots show the result of Friedman’s test to assess phase modulation.

We first describe cutaneous reflexes in the ipsilateral SOL (**Fig. 2**), a uni-articular ankle extensor (plantar flexor) that activated before hindlimb contact and remained active for approximately 80% of the stance period. When SOL was active, stimulating the SP nerve during quadrupedal locomotion evoked short-latency inhibition, or N1 responses in 100% of nerves stimulated (n = 11/11, **Table 1**) and in 58% of bins in the cycle (n = 64/110). N1 responses are only visible when a muscle is active. These results are consistent with previous findings from intact cats during quadrupedal locomotion (Bretzner and Drew 2005; Duysens and Loeb 1980; Forssberg 1979; Forssberg et al. 1977; Frigon et al. 2009; Frigon and Rossignol 2007, 2008b; Hurteau et al. 2017, 2018; Loeb 1993; Pratt et al. 1991). During hindlimb-only locomotion, N1 responses were also present in all nerves stimulated (n = 11/11, 100%) and in 59% of bins in the cycle (n = 65/110). We found no significant differences in N1 response occurrence between locomotor conditions (P = 0.891). The N1 responses were significantly modulated by phase in both locomotor conditions (Quadrupedal, P = 1.03^e-12^; Hindlimb-only, P = 1.61^e-12^). Studies have shown that N1 responses are followed by mid-(P2) or longer-latency (P3) excitatory responses (Abraham et al. 1985; Duysens and Loeb 1980; Frigon et al. 2009; Frigon and Rossignol 2008a; Loeb 1993). Considering that the duration of the inhibitory response influences the appearance of the excitatory response, we combined the mid and long-latency responses (P2/P3) together. In this study, we observed P2/P3 responses in the ipsilateral SOL with 73% of nerves stimulated (n = 8/11) during quadrupedal locomotion but only in 27% of nerves stimulated during hindlimb-only locomotion (n = 3/11). We also found significantly more P2/P3 responses during quadrupedal locomotion (n = 49/80 bins or 61%) compared to hindlimb-only locomotion (n = 10/30 bins or 33%) (P = 2.93^e-9^). During quadrupedal locomotion, P2/P3 responses in the ipsilateral SOL were significantly modulated by phase for pooled data (P = 0.007), but not during hindlimb-only locomotion (P = 0.161). During quadrupedal locomotion, we also observed short-latency excitation, or P1 responses in 36% of nerves stimulated (n = 4/11) and 38% of bins in the cycle (n = 15/40), mainly when the ipsilateral SOL was inactive, consistent with previous findings (Frigon et al. 2009). During hindlimb-only locomotion, P1 responses were present in 27% of nerves stimulated (n = 3/11) and 33% of bins in the cycle (n = 10/30), with no significant differences between locomotor conditions (P = 0.288). The P1 responses evoked by ipsilateral SP nerve stimulation were significantly modulated by phase for pooled data in both locomotor tasks (Quadrupedal, P = 2.22^e-3^; Hindlimb-only, P = 0.005).

The ipsilateral VL and BFA (**Fig. 2**), knee and hip extensor muscles, respectively, became active around foot contact and throughout the majority of the stance phase. As for the ipsilateral SOL, when VL and BFA are active, stimulating cutaneous or mixed nerves of the foot during quadrupedal locomotion in intact cats evoked N1 responses followed by P2/P3 responses, as shown previously (Abraham et al. 1985; Duysens and Loeb 1980; Frigon and Rossignol 2008b; Loeb 1993). In the present study, N1 responses were present in 100% of nerves stimulated for the ipsilateral VL in both locomotor conditions (Quadrupedal, n = 9/9; Hindlimb-only, n = 9/9; **Table 1**). For the ipsilateral BFA, N1 responses were present in more nerves stimulated during quadrupedal locomotion (n = 11/11) compared to hindlimb-only locomotion (n = 8/11). For both muscles, we observed a similar number of bins with N1 responses during quadrupedal locomotion (VL, n = 59/90 bins or 66%; BFA, n = 66/110 bins or 60%) and hindlimb-only locomotion (VL, n = 63/90 bins or 70%; BFA, n = 52/80 bins or 65%), with no significant difference in reflex occurrence between locomotor conditions (VL, P = 0.587; BFA, P = 0.058). In both tasks, N1 responses were significantly modulated by phase for VL (Quadrupedal, P = 4.84^e-8^; Hindlimb-only, P = 7.41^e-7^) and BFA (Quadrupedal, P = 4.78^e-11^; Hindlimb-only, P = 2.00^e-5^). For the ipsilateral VL, we observed P2/P3 responses in 67% of nerves stimulated (n = 6/9) during quadrupedal locomotion and in 56% of nerves stimulated during hindlimb-only locomotion (n = 5/9). The number of bins in the cycle with P2/P3 responses was similar during quadrupedal locomotion (n = 25/60 bins or 42%) and hindlimb-only locomotion (n = 19/50 bins or 38%), with no significant differences between locomotor conditions (P = 0.312). The P2/P3 responses in the ipsilateral VL were not significantly modulated by phase for pooled data (Quadrupedal, P = 0.274; Hindlimb-only, P = 0.061). For the ipsilateral BFA, we observed P2/P3 responses in 91% of nerves stimulated (n = 10/11) during quadrupedal locomotion but only in 18% of nerves stimulated during hindlimb-only locomotion (n = 2/11). P2/P3 responses were more frequent during quadrupedal locomotion (n = 55/100 bins or 55%) compared to hindlimb-only locomotion (n = 7/20 bins or 35%) (P = 6.32^e-13^). During quadrupedal locomotion, P2/P3 responses in the ipsilateral BFA were significantly modulated by phase for pooled data (P = 0.002). During the inactive phase, we found P1 responses in a single cat in both locomotor conditions.

Next, we evaluated the modulation of cutaneous reflexes in ipsilateral muscles that flex the hip (SRT), knee (ST) and ankle (TA) joints, although the SRT and ST also extend the knee and hip, respectively (**Fig. 3**). The SRT burst covered most of the hindlimb swing phase during quadrupedal locomotion in intact cats. During hindlimb-only locomotion the SRT also displayed a prominent second burst during the stance phase, as shown previously (Harnie et al. 2022; Klishko et al. 2021; Smith et al. 1998). In the literature, stimulating the SP nerve during quadrupedal locomotion in intact cats evokes N1 responses followed by P2 responses when SRT is active and mainly P2 responses when inactive (Hurteau et al. 2018). Here, we observed N1 responses in 50% of nerves stimulated (n = 4/8, **Table 1**) and in 68% of bins (n = 27/40) during quadrupedal locomotion in SRT. Interestingly, N1 responses were present in 100% of nerves stimulated (n = 8/8) during hindlimb-only locomotion and in 74% of bins (n = 59/80 bins), a significant increase in response occurrence compared to quadrupedal locomotion (P = 1.0^e-4^). N1 responses in the ipsilateral SRT were not significantly modulated by phase for pooled data during quadrupedal locomotion (P = 0.146) and hindlimb-only locomotion (P = 0.116). We also observed P2 responses in both locomotor conditions. During quadrupedal locomotion, we found P2 responses in 25% of nerves stimulated (n = 2/8) and in 35% of bins (n = 7/20). These excitatory responses appeared at the time of liftoff and just before the onset of muscle activation. During hindlimb-only locomotion, we found P2 responses in 50% of nerves stimulated (n = 4/8) and in 38% of bins (n = 15/40), with no significant differences in occurrence between conditions (P = 0.072). P2 responses evoked during hindlimb-only locomotion were not significantly modulated by phase for pooled data (P = 0.056) and we did not have sufficient data (n = 2) for quadrupedal locomotion.

**Figure 3.**
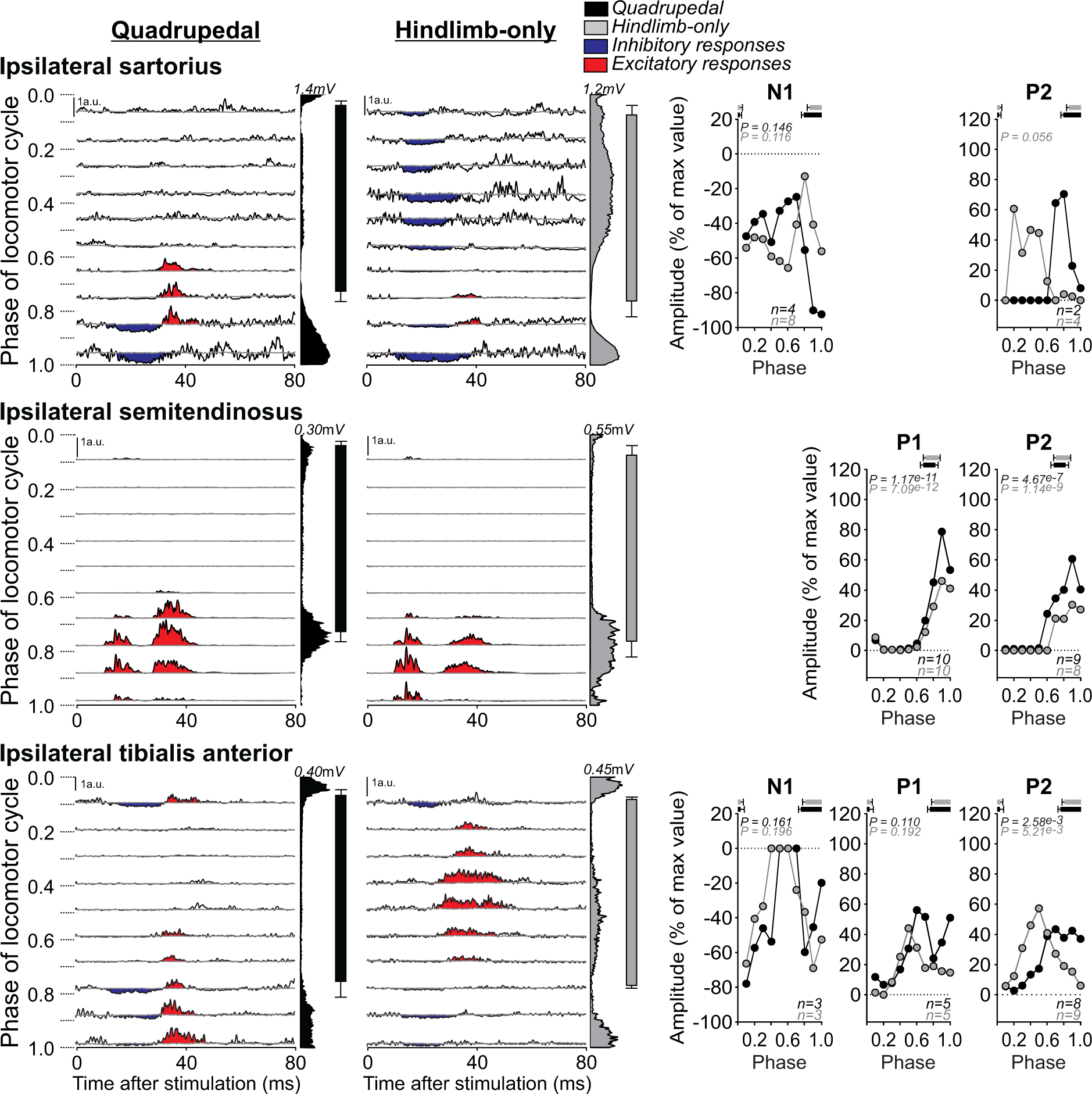
Phase- and condition-dependent modulation of cutaneous reflexes in ipsilateral muscles that flex the hip, knee and ankle joints. SP nerve stimulation evoked responses in the ipsilateral sartorius (n = 8 nerves in 6 cats), semitendinosus (n = 10 nerves in 6 cats) and tibialis anterior (n = 10 nerves in 6 cats) during quadrupedal and hindlimb-only locomotion. Each panel represents averaged waveforms in the 10 bins of the cycle with a post-stimulation window of 80 ms in a single cat (sartorius and semitendinosus, Cat 1; tibialis anterior, Cat 3). Black traces represent averaged cycles that received a stimulation (i.e., reflex responses, n = 4 – 22 stimuli per bin). Gray traces represent averaged cycles with no stimuli. Scale bars are shown in arbitrary units (a.u.) for both locomotor conditions. Aligned vertically is the average rectified EMG and the stance phase aligned to ipsilateral soleus muscle onset. Scatter plots represent reflex amplitudes (mean) in each bin of the cycle expressed as a percentage of the maximal response found in one of the bins for each nerve stimulated during quadrupedal (black circles) and hindlimb-only locomotion (gray circles). Short-latency (P1, N1) and mid-latency (P2) responses are shown separately. Horizontal bars at the top of the scatter plots represent the period of muscle activity for each locomotor condition (n = 9-14 control cycles) for pooled data. P values at the top of the scatter plots show the result of Friedman’s test to assess phase modulation.

The ST is a knee flexor and hip extensor muscle that displayed a short burst at swing onset in both locomotor conditions and a prominent second burst during early stance in hindlimb-only locomotion at the same speed (Harnie et al. 2022). In the present study, we observed P1 responses in the ipsilateral ST in 100% of nerves stimulated (n = 10/10, **Table 1**) and in 55% of bins (n = 55/100) during quadrupedal locomotion. During hindlimb-only locomotion, we also found P1 responses in 100% of nerves stimulated (n = 10/10) and in 50% of bins (n = 50/100), with no significant difference in occurrence between locomotor conditions (P = 0.500). In both tasks, P1 responses were significantly modulated by phase (Quadrupedal, P = 1.17^e-11^; Hindlimb-only, P = 7.09^e-12^), with prominent responses during the activity of the ipsilateral ST and throughout swing, whereas they were generally suppressed during stance. P1 responses were generally followed by P2 responses, as shown by several studies during quadrupedal locomotion in intact cats (Bretzner and Drew 2005; Frigon et al. 2009; Hurteau et al. 2018; Hurteau and Frigon 2018; Pratt et al. 1991). Here, we observed P2 responses in the ipsilateral ST in 90% of nerves stimulated (n = 9/10) and 46% of bins (n = 41/90) during quadrupedal locomotion. During hindlimb-only locomotion, we found P2 responses in 80% of nerves stimulated (n = 8/10) and 45% of bins (n = 36/80), with no significant difference in occurrence between locomotor conditions (P = 0.113). In both tasks, P2 responses were significantly modulated by phase (Quadrupedal, P = 4.67^e-7^; Hindlimb-only, P = 1.14^e-9^).

The TA is a uni-articular ankle flexor, with a main burst covering most of the swing phase during quadrupedal locomotion. However, in two cats a second burst was also observed during the stance phase. We found the same pattern of muscle activation during hindlimb-only locomotion, with an additional cat displaying a second burst during stance. Here, stimulating the SP nerve evoked responses in the ipsilateral TA that differed based on cats and nerves stimulated (**Fig. 3**). We found N1 responses at the beginning and end of the cycle in 30% of nerves stimulated in both locomotor conditions (n = 3/10, **Table 1**). We observed no significant difference in N1 response occurrence between locomotor conditions (P = 0.686). N1 responses were not significantly modulated by phase (Quadrupedal, p = 0.161; Hindlimb-only, P = 0.196). We also found P1 responses in 50% of nerves stimulated (n = 5/10) and 72% of bins (n = 36/50) during quadrupedal locomotion and in 50% of nerves stimulated (n = 5/10) and 48% of bins (n = 24/50) during hindlimb-only locomotion, with no significant difference in response occurrence between locomotor conditions (P = 0.098). In both tasks, P1 responses were not significantly modulated by phase (Quadrupedal, P = 0.110; Hindlimb-only, P = 0.192). We also found P2 responses in 80% of nerves stimulated (n = 8/10) during quadrupedal locomotion and in 54% of bins (n = 43/80), mainly during swing. During hindlimb-only locomotion, we observed P2 responses in 90% of nerves stimulated (n = 9/10) and 58% of bins (n = 52/90), mainly during stance, with no significant difference in P2 response occurrence between locomotor condition (P = 0.221). In both tasks, P2 responses were significantly modulated by phase (Quadrupedal, P = 2.58^e-3^; Hindlimb-only, P = 5.21^e-3^).

### Cutaneous reflex responses in contralateral muscles

Responses evoked by electrical stimuli of the foot dorsum in ipsilateral limb muscles lift the limb away from and over the simulated contact of the foot dorsum during swing, while responses in contralateral limb muscles mainly reinforce weight support (Buford and Smith 1993; Duysens and Loeb 1980; Forssberg 1979; Forssberg et al. 1977; Hurteau et al. 2017, 2018; Hurteau and Frigon 2018; Prochazka et al. 1978). Here, we investigated cutaneous reflexes in four muscles (SOL, VL, BFA and SRT) contralateral to the stimulation (**Fig. 4 and Table 1**). We did not include responses in contralateral ST and TA because we only found them in one cat.

**Figure 4.**
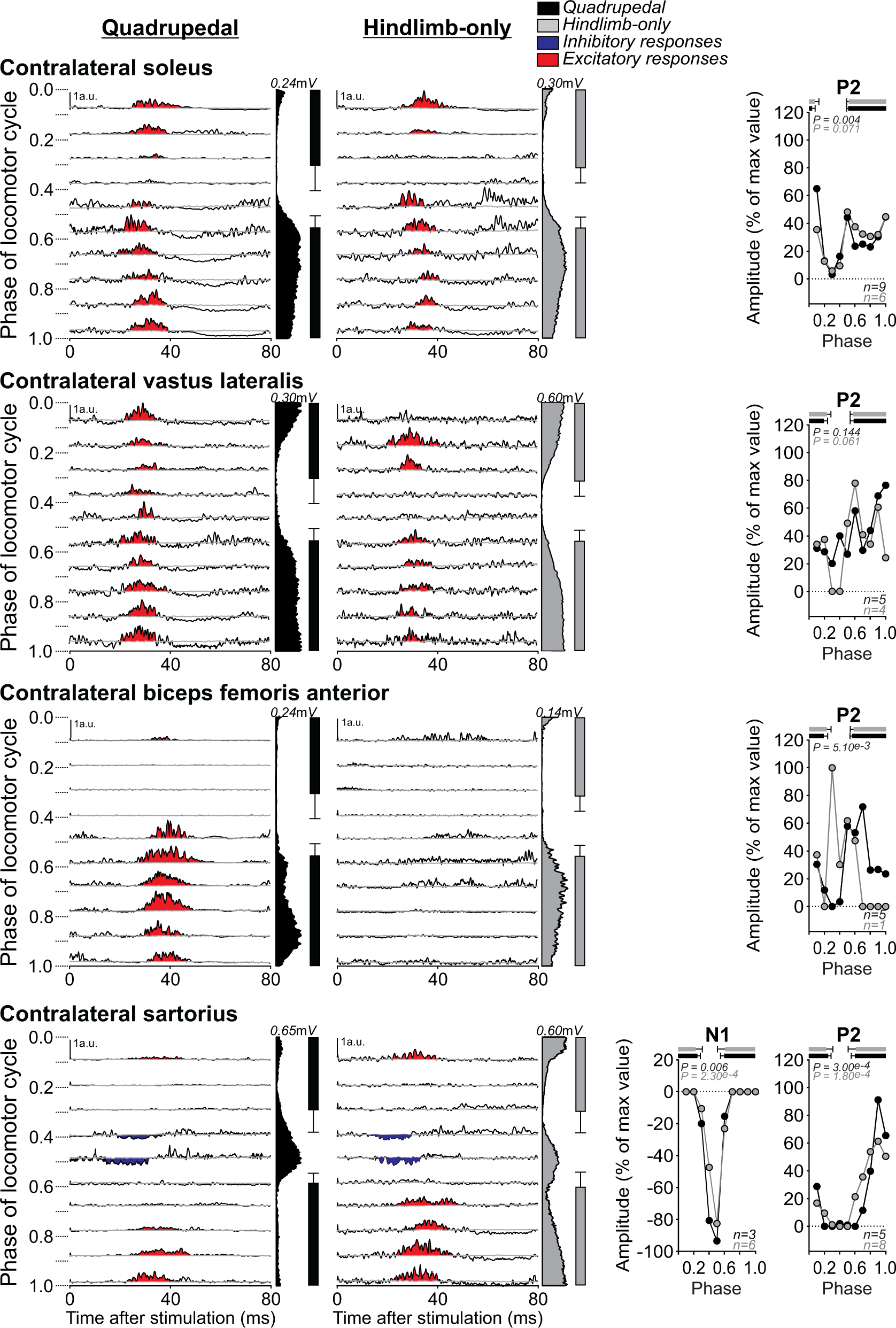
Phase- and condition-dependent modulation of cutaneous reflexes in contralateral muscles. SP nerve stimulation evoked responses in the contralateral soleus (n = 10 nerves in 6 cats), vastus lateral (n = 6 nerves in 5 cats), biceps femoris anterior (n = 6 nerves in 5 cats) and sartorius (n = 8 nerves in 6 cats) during quadrupedal and hindlimb-only locomotion. Each panel represents averaged waveforms in the 10 bins of the cycle with a post-stimulation window of 80 ms in a single cat (cat 1 for the three muscles). Black traces represent averaged cycles that received a stimulation (i.e., reflex responses, n = 4–22 stimuli per bin). Gray traces represent averaged cycles with no stimuli. Scale bars are shown in arbitrary units (a.u.) for both locomotor conditions. Aligned vertically is the average rectified EMG and the stance phase aligned to ipsilateral soleus muscle onset. Scatter plots represent reflex amplitudes (mean) in each bin of the cycle expressed as a percentage of the maximal response found in one of the bins for each nerve stimulated during quadrupedal (black circles) and hindlimb-only locomotion (gray circles). Horizontal bars at the top of the scatter plots represent the period of muscle activity for each locomotor condition (n = 9-15 control cycles) for pooled data. P values at the top of the scatter plots show the result of Friedman’s test to assess phase modulation.

In the contralateral SOL (**Fig. 4**), we observed P2 responses in 82% of nerves stimulated (n = 9/11) and 64% of bins (n = 58/90) during quadrupedal locomotion. During hindlimb-only locomotion, we observed P2 responses in 55% of nerves stimulated (n = 6/11) and 72% of bins (n = 43/60), a significant reduction in response occurrence compared to quadrupedal locomotion (P = 0.042). Although we observed the same overall profile of phase-dependent modulation, P2 responses were significantly modulated by phase for pooled data during quadrupedal (P = 0.004) but not hindlimb-only (P = 0.071) locomotion.

In the contralateral VL and BFA (**Fig. 4**), we observed P2 responses during the muscles’ active periods in both locomotor conditions. For contralateral VL, we found P2 responses in 56% of nerves stimulated (n = 5/9, **Table 1**) and 74% of bins (n = 37/50) during quadrupedal locomotion and in 44% of nerves stimulated (n = 4/9) and 68% of bins (n = 27/40) during hindlimb-only locomotion, with no significant difference in P2 response occurrence between conditions (P = 0.138). In both tasks, P2 responses in the contralateral VL were not significantly modulated by phase (Quadrupedal, P = 0.144; Hindlimb-only, P = 0.061). For the contralateral BFA, we found P2 responses in 45% of nerves stimulated (n = 5/11) and 74% of bins (n = 37/50) during quadrupedal locomotion but during hindlimb-only locomotion, we observed P2 responses in only 9% of nerves stimulated (n = 1/11) and 50% of bins in the cycle (n = 5/10), a significant reduction in response occurrence compared to quadrupedal locomotion (P = 4.03^e-8^). During quadrupedal locomotion, P2 responses were significantly modulated by phase (P = 5.10^e-3^).

In the contralateral SRT (**Fig. 4**), we observed N1 responses during the swing phase in 38% of nerves stimulated (n = 3/8; **Table 1**) and 27% of bins (n = 8/30) during quadrupedal locomotion and in 75% of nerves stimulated (n = 6/8) and 22% of bins (n = 13/60) during hindlimb-only locomotion, with no significant difference in response occurrence between conditions (P = 0.251). In both tasks, N1 responses were significantly modulated by phase (Quadrupedal, P = 0.006; Hindlimb-only, P = 2.30^e-4^). Stimulating the SP nerve evoked P2 responses in 63% of nerves stimulated (n = 5/8) and 44% of bins (n = 22/50) during quadrupedal locomotion and in 100% of nerves stimulated (n = 8/8) and 48% of bins (n = 38/80) during hindlimb-only locomotion, a significant increase in response occurrence compared to quadrupedal locomotion (P = 0.015). In both locomotor conditions, P2 responses were significantly phase modulated (Quadrupedal, P = 3.00^e-4^; Hindlimb-only, P = 1.80^e-4^).

The symbols (arrows and equal signs) in **Table 1** summarize response occurrence during quadrupedal and hindlimb-only locomotion for the different ipsilateral and contralateral muscles. We can summarize response occurrence as follows. N1 and P1 response occurrences were similar in both conditions, with the exception of the ipsilateral SRT where it was greater during hindlimb-only locomotion. P2/P3 response occurrences were reduced during hindlimb-only locomotion in ipsilateral SOL, ipsilateral BFA, contralateral SOL and contralateral BFA but increased in contralateral SRT. In the ipsilateral VL, SRT, ST and TA as well as contralateral VL, P2/P3 response occurrences were similar in both conditions.

### Hindlimb-only locomotion reduces cutaneous reflex modulation in the ipsilateral semitendinosus

To compare reflex responses evoked in the two conditions, we measured a modulation index by subtracting the smallest response from the largest response observed in the 10 bins (**Fig. 5**) (Hurteau et al. 2017, 2018). In the ipsilateral ST muscle, we observed a significantly smaller reflex modulation during hindlimb-only locomotion compared to quadrupedal locomotion for P1 (P = 0.022; 41% difference) and P2 (P = 0.017; 55% difference) responses. In the other muscles where we had a sufficient number of excitatory reflex responses for statistical analysis, the modulation index did not significantly differ between the two locomotor conditions. This was the case for P1 (P = 0.686) and P2 (P = 0.779) responses in the ipsilateral TA, P2 responses in the contralateral SOL (P = 0.249) and P2 responses in the contralateral SRT (P = 0.080).

**Figure 5.**
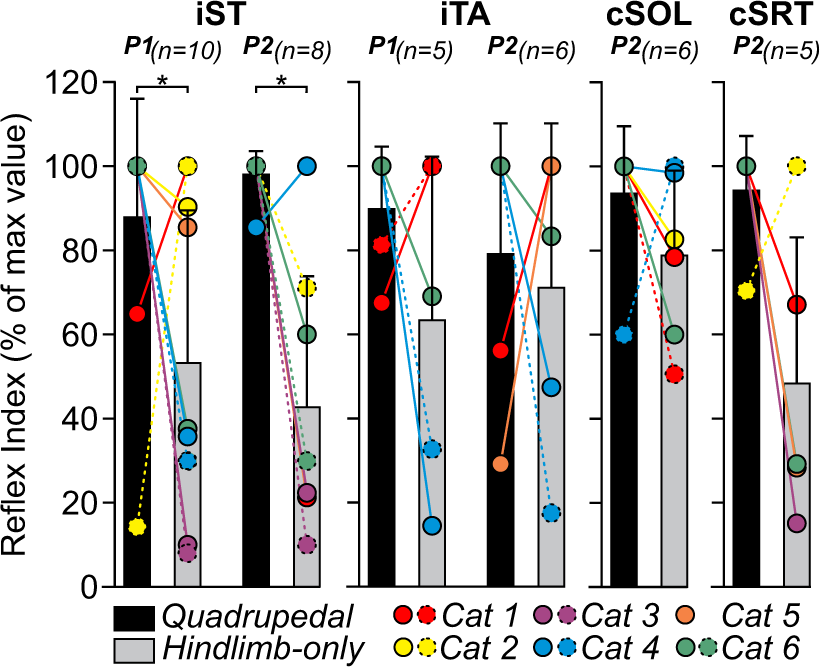
Cutaneous reflex modulation across locomotor conditions for pooled data. Each bar represents the mean reflex index ± standard deviation for pooled data of responses evoked in the ipsilateral semitendinosus (iST) and tibialis anterior (iTA) as well as the contralateral soleus (cSOL) and sartorius (cSRT). The coloured circles represent the nerves stimulated in the 6 cats. The solid and dotted lines of the circles are for the right and left SP nerves, respectively. Asterisks indicate significant differences between conditions (Wilcoxon): *p<0.05.

### Cutaneous reflex responses in intact and spinal cats

In previous sections, we described cutaneous reflexes during quadrupedal and hindlimb-only locomotion in intact cats. In this section, we compare cutaneous reflexes evoked in two cats (n = 4 nerves stimulated) in the two intact conditions and during hindlimb-only locomotion after spinal transection. We did this to determine if changes in cutaneous reflex response in spinal cats are due to the lesion or to the absence of forelimb movements. **Figure 6** shows cutaneous reflex responses in three ipsilateral muscles at their periods of mid-activity and mid-inactivity.

**Figure 6.**
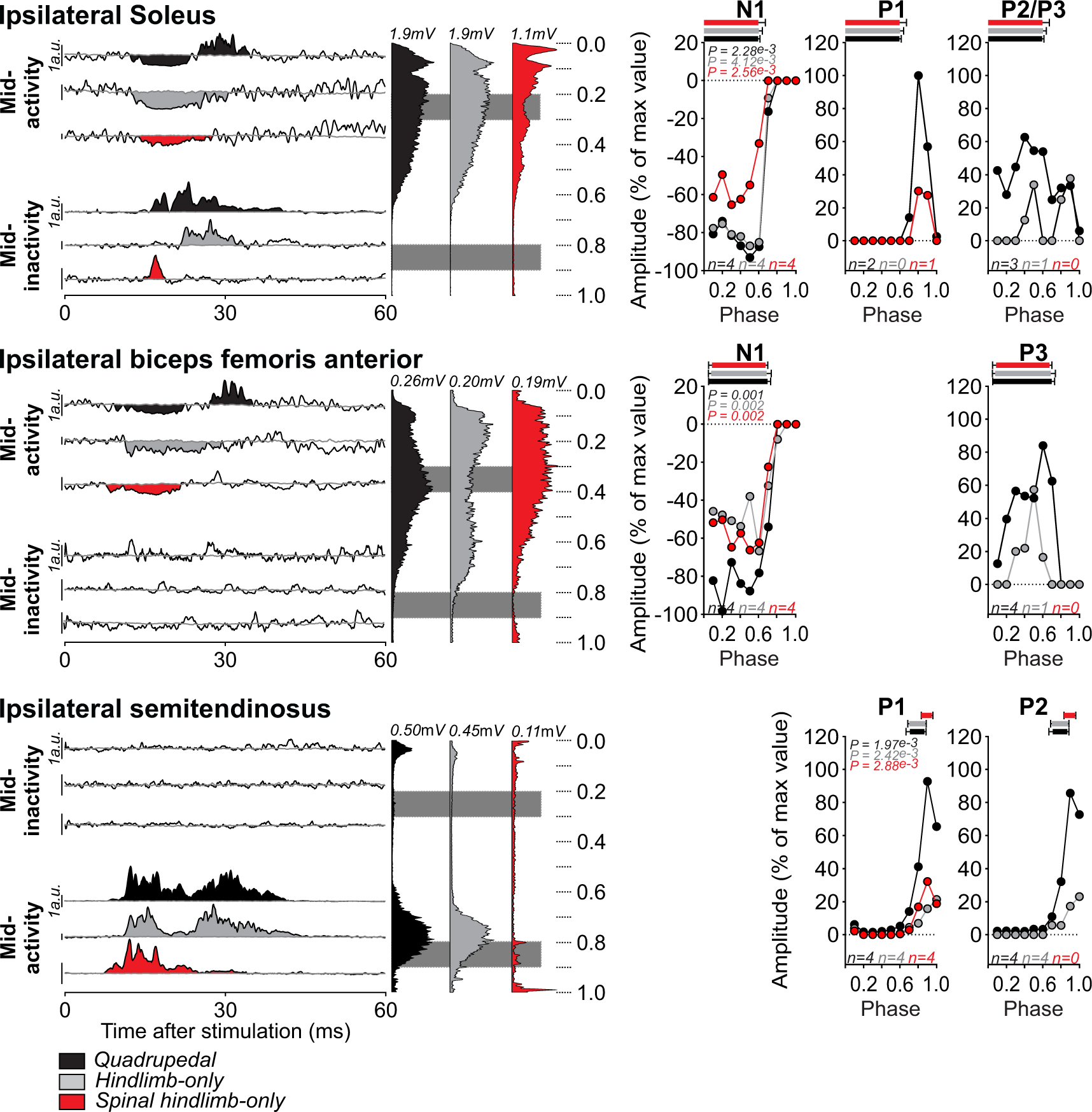
Phase- and condition-dependent modulation of cutaneous reflexes in intact and spinal cats. SP nerve stimulation evoked responses in the ipsilateral soleus (n = 4 nerves in 2 cats), biceps femoris anterior (n = 3 nerves in 2 cats) and semitendinosus (n = 4 nerves in 2 cats) in the intact state during quadrupedal and hindlimb-only locomotion and after spinal transection in the hindlimb-only condition. Each panel represents averaged waveforms at mid-activity and mid-inactivity of the muscle’s main burst with a post-stimulation window of 80 ms from a single cat (soleus, Cat 3; biceps femoris anterior, Cat 6; semitendinosus, Cat 3). Black traces represent averaged cycles that received a stimulation (i.e., reflex responses, n = 7–18 stimuli per bin). Gray traces represent averaged cycles with no stimuli. Scale bars are shown in arbitrary units (a.u.) for both conditions of locomotion. Aligned vertically is the average rectified EMG aligned to ipsilateral soleus muscle onset. The grey areas correspond to the bins taken in periods of mid activity and mid inactivity of the muscle. Scatter plots represent reflex amplitudes (mean) in each bin of the cycle expressed as a percentage of the maximal response found in one of the bins for each nerve stimulated between quadrupedal (black circles), hindlimb-only locomotion (gray circles) in the intact state and hindlimb-only condition in the spinal sate (red circles). Short-latency (P1, N1) and mid/longer-latency (P2/P3) responses are shown separately. Horizontal bars at the top of the scatter plots represent the period of muscle activity for each locomotor condition (n = 7-15 control cycles) for pooled data. P values at the top of the scatter plots show the result of Friedman’s test to assess phase modulation.

In the ipsilateral SOL, we observed N1 responses in 100% of nerves stimulated (n = 4/4) in both intact locomotor conditions and during hindlimb-only locomotion in the spinal state. In all three conditions, N1 responses were significantly modulated by phase (Quadrupedal, P = 2.28^e-3^; Hindlimb-only, P = 4.12^e-3^; Spinal hindlimb-only, P = 2.56^e-3^). While mid- or longer-latency excitatory responses (P2/P3) following the N1 responses were found in 75% of nerves stimulated during intact quadrupedal locomotion (n = 3/4), P2/P3 responses were only present in 25% of nerves stimulated during intact hindlimb-only locomotion (n = 1/4) and not at all in the spinal state (n = 0/4). During the SOL’s inactive phase, P1 responses were observed in 50% (n = 2/4), 0% (n = 0/4) and 25% (n = 1/4) of nerves stimulated during quadrupedal and hindlimb-only locomotion in the intact state and during hindlimb-only locomotion in the spinal state, respectively.

During the active period of the ipsilateral BFA, we observed N1 responses in 100% of nerves stimulated (n = 4/4) in both intact conditions and during hindlimb-only locomotion in the spinal state. In all three conditions, N1 responses were significantly modulated by phase (Quadrupedal, P = 0.001; Hindlimb-only, P = 0.002; Spinal hindlimb-only, P = 0.002). We observed longer-latency excitatory responses (P3) in 100% of nerves stimulated during quadrupedal locomotion (n = 4/4) but in only 25% of nerves stimulated during intact hindlimb-only locomotion (n = 1/4) and not at all in the spinal state (n = 0/4). During the ipsilateral BFA’s inactive phase, we did not observe P1 or P2 responses in all three locomotor conditions.

During the active period of the ipsilateral ST, we observed P1 responses in 100% of nerves stimulated (n = 4/4) in both intact conditions and during hindlimb-only locomotion in the spinal state. In all three conditions, P1 responses were significantly modulated by phase (Quadrupedal, P = 1.97^e-3^; Hindlimb-only, P = 2.42^e-3^; Spinal hindlimb-only, P = 2.88^e-3^). While the P2 responses following the P1 responses were observed in 100% of nerves stimulated (n = 4/4) in both intact conditions, we found no P2 responses in the spinal state.

## DISCUSSION

In the present study, we investigated task-dependent modulation of hindlimb cutaneous reflexes by comparing responses evoked during quadrupedal and hindlimb-only locomotion to determine the contribution of forelimb movements to the modulation of hindlimb reflex pathways. Consistent with our hypothesis, we found differences in reflex responses between the two conditions. This is not surprising because forelimb and hindlimb movements must be coordinated during quadrupedal locomotion. Depending on the muscle, not all nerve stimulations elicited a response during quadrupedal or hindlimb-only locomotion. We found that the occurrence of short-latency inhibitory or excitatory responses was similar in most muscles in both conditions, with the exception of the ipsilateral SRT, a hip flexor/knee extensor, where short-latency inhibitory responses were more frequent during hindlimb-only locomotion. Thus, the absence of forelimb movements has little effect on short-latency reflex pathways. On the other hand, the occurrence of longer-latency excitatory responses decreased in some extensor muscles bilaterally (SOL and BFA) during hindlimb-only locomotion, with the exception of the contralateral SRT where it increased. These results and those in spinal cats suggest that the absence of forelimb movements in intact cats suppresses spino-bulbo-spinal pathways (Shimamura et al. 1967) that contribute to longer-latency excitatory responses in extensor muscles (i.e. those mostly active during stance) while having little to no effect on short-latency responses. In the next sections, we discuss the contribution of forelimb movements in the modulation of hindlimb reflex responses/pathways and its functional significance for locomotor control.

### Factors influencing reflex responses during quadrupedal and hindlimb-only locomotion

Several factors can influence reflex responses. One is the background level of EMG, or automatic gain compensation (Burke 2016; Capaday and Stein 1987; Matthews 1986). In general, larger and/or more frequent responses are evoked when a muscle is active because more motoneurons and premotor interneurons are firing or close to threshold when the incoming sensory inputs arrive and negative/inhibitory responses require a certain baseline level of EMG activity to be visible and measured. Thus, if a muscle stays active longer, it increases the probability of evoking or observing a response, particularly an inhibitory response, and conversely, fewer responses are observed if EMG activity is weaker or shorter, particularly excitatory responses. A greater occurrence of N1 responses in the ipsilateral SRT and P2/P3 responses in the contralateral SRT during hindlimb-only locomotion can be explained by the prominent second burst in this muscle during the stance phase (**Fig. 1A**). The stance-related activity of the SRT muscle during hindlimb-only locomotion was shown by others (Harnie et al. 2022; de Leon et al. 1998). However, the reduction of P2/P3 responses in SOL and BFA, ipsilateral and contralateral to the stimulation, during hindlimb-only locomotion cannot be explained by shorter periods of activity because the stance-related BFA burst occupied a greater proportion of the cycle compared to quadrupedal locomotion in intact cats while there was no difference in SOL (**Fig. 1C**). The mean amplitude of SOL was significantly reduced during hindlimb-only locomotion, however (**Fig. 1D**). In hindlimb-only locomotion, the forelimbs are slightly elevated on a stationary platform. This likely produces a caudal shift of the body’s center of mass, potentially increasing loading on the hindlimbs and the amplitude of EMG activity of extensor muscles. We observed a significant increase in the EMG amplitude of VL, a knee extensor, during hindlimb-only locomotion compared to quadrupedal locomotion but a decrease in SOL, an ankle extensor, and no difference in BFA, a hip extensor (**Fig. 1**). The occurrence of short-latency inhibitory reflex responses in the ipsilateral SOL, VL and BFA was similar in quadrupedal and hindlimb-only locomotion. Thus, the background level of EMG did not necessarily translate into greater or fewer reflex responses, as shown by others in cats (Hurteau et al. 2017, 2018; Mari et al. 2023; Quevedo et al. 2005a; Zehr et al. 1997) or humans (Zehr et al. 1997).

Another potential factor affecting cutaneous reflex responses during locomotion is the symmetry between the left and right sides, particularly its spatial component, as shown in intact and spinal cats (Hurteau et al. 2017; Hurteau and Frigon 2018; Mari et al. 2023). However, this is unlikely a factor in reflex modulation/response occurrence in the present study because we found no significant differences in temporal or spatial left-right asymmetry between quadrupedal and hindlimb-only locomotion in intact cats (**Fig. 1E**). This is not surprising because tied-belt locomotion at 0.4 m/s is inherently symmetric, with a strict out-of-phase alternation of the hindlimbs (Dambreville et al. 2015).

### Functional considerations

Electrically stimulating the superficial peroneal nerve mimics a mechanical contact to the dorsum of the hindpaw, generating a stumbling corrective reaction during swing and a stumbling preventive reaction during stance. In cats, the main muscles responsible for the stumbling corrective reaction are muscles that flex the knee and extend the hip, such as ST and posterior biceps femoris, which move the limb away and vertically from the contact point during swing (Buford and Smith 1993; Forssberg et al. 1977). In the present study, we found a significant reduction in the depth of reflex modulation of short- and mid-latency excitatory responses in the ipsilateral ST muscle during hindlimb-only locomotion compared to quadrupedal locomotion (**Fig. 5**). This would reduce knee flexion/hip extension during the swing phase following contact of the foot dorsum during hindlimb-only locomotion. We also observed fewer longer-latency excitatory responses in extensor muscles (SOL and BFA) of the contralateral hindlimb during the swing phase of hindlimb-only locomotion. The less frequent responses in contralateral muscles might be because the ipsilateral hindlimb flexes less (smaller excitatory responses in the ipsilateral ST), requiring less contralateral support, and because the body is in a more stable position during hindlimb-only locomotion. In quadrupedal locomotion at 0.4 m/s on a treadmill, the pattern consists of alternating periods of double and triple support, where two and three limbs are contacting the surface, respectively, with some brief and infrequent periods of quadruple support, where four limbs contact the surface, which replace diagonal support periods (Frigon et al. 2014). In hindlimb-only locomotion, as the forelimbs always contact the surface, the pattern alternates between periods of triple and quadruple support, which offers greater stability.

With stimulation during ipsilateral stance (i.e. the stumbling preventive reaction), we found more frequent inhibitory responses in the ipsilateral SRT, a hip flexor/knee extensor, and fewer longer-latency excitatory responses in some ipsilateral extensor muscles (SOL and BFA) during hindlimb-only locomotion. As the SRT shows a prominent burst in the stance phase of hindlimb-only locomotion and because it flexes the hip, it makes functional sense that it receives inhibition during stance to prevent the limb from flexing and potentially destabilizing the pattern. Fewer excitatory responses in extensor muscles might be because the animal is in a more stable position and the longer-latency responses, which are likely mediated by supraspinal loops/pathways, are not required. Longer-latency reflex responses are thought to allow supraspinal structures to integrate somatosensory information and modify limb trajectory and posture according to phase and state (Pruszynski and Scott 2012).

### Spinal versus supraspinal contributions

One way to investigate the relative contribution of supraspinal versus spinal mechanisms in evoking and modulating hindlimb reflexes is to compare responses before and after spinal transection. After thoracic spinal transection, all communication between the brain/cervical cord and lumbosacral circuits controlling hindlimb movements are permanently disrupted. Although the stumbling corrective reaction is observed in spinal cats and stimulating the SP nerve evokes phase-dependent reflex responses, mid- and long-latency responses in hindlimb muscles are generally reduced or abolished (**Fig. 6**), consistent with an important supraspinal contribution to these responses (Forssberg et al. 1975, 1977; Frigon and Rossignol 2008b; Hurteau et al. 2017; Hurteau and Frigon 2018; LaBella et al. 1992). In the present study, we showed that the absence of forelimbs movements has little effect on short-latency responses but that longer-latency responses are suppressed in intact cats. This reinforces the notion that short-latency reflexes are mainly mediated and modulated by spinal circuits/mechanisms while longer-latency responses require contributions from supraspinal structures and/or sensorimotor circuits controlling the forelimbs in the cervical cord. As the forelimbs do not need to be coordinated with the hindlimbs during hindlimb-only locomotion, the descending pathways from the brain and cervical cord that contribute to longer-latency responses in hindlimb muscles are suppressed.

We currently do not know where longer-latency reflex pathways are gated. **Figure 7** provides a plausible scenario that remains speculative. In quadrupedal locomotion, primary afferents from the SP nerve make contact with spinal interneurons that generate short-latency inhibitory or excitatory responses in ipsilateral muscles as well as with interneurons with crossed projections that generate longer latency excitatory responses in contralateral muscles (**Fig. 7A**). Afferents inputs from the SP nerve also contact neurons that ascend via propriospinal or dorsal column tracts to the cervical cord and supraspinal structures. These ascending inputs make contact with neurons in the brainstem that then send descending inputs to the lumbar cord and contribute to longer-latency reflex responses in ipsilateral and/or contralateral muscles (Shimamura and Aoki 1969). The absence of forelimb movements during hindlimb-only locomotion suppresses ascending inputs to supraspinal structures and/or their descending influences (**Fig. 7B**). These responses can be gated at lumbar, cervical and/or supraspinal levels. After thoracic spinal transection, ascending inputs no longer reach the cervical cord and supraspinal structures no longer contribute to longer-latency reflex responses (**Fig. 7C**).

**Figure 7.**
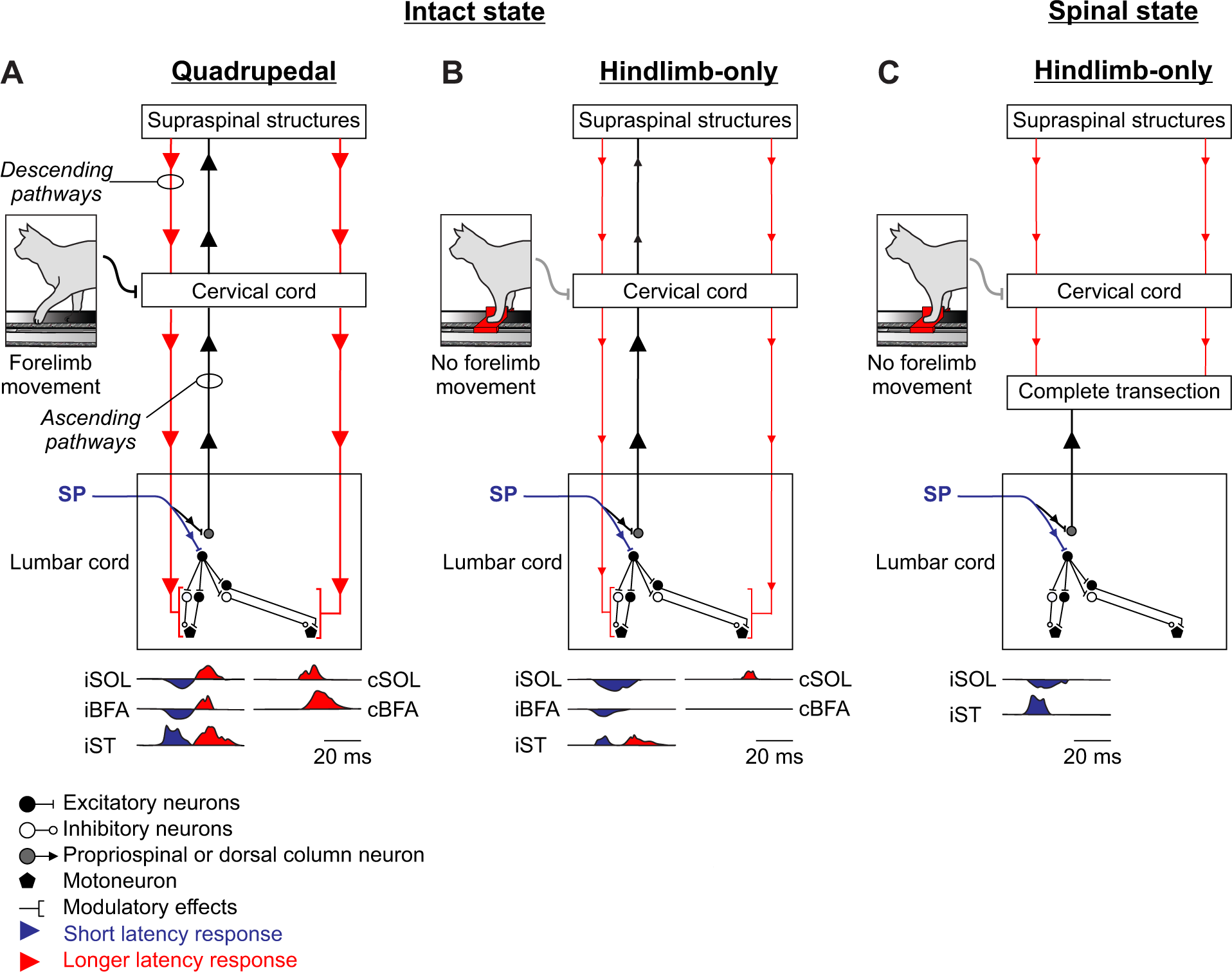
Spinal and supraspinal contributions to short- and longer-latency reflex responses during quadrupedal and hindlimb-only locomotion. The figure shows possible spinal and supraspinal contributions to reflex responses from superficial peroneal (SP) afferents in the intact and spinal states during quadrupedal and hindlimb-only locomotion. **A)** During intact quadrupedal locomotion, SP afferents activate lumbar circuits that elicit short-latency reflex responses in the ipsilateral hindlimb, and in the contralateral hindlimb via crossed projections. Afferent inputs also activate neurons that project to circuits of the cervical cord and brain, which then project back to lumbar circuits to evoke longer-latency responses. **B)** During hindlimb-only locomotion in the intact state, SP afferents make the same synaptic contacts but the absence of forelimb movements suppresses descending signals from the brain and cervical cord that contribute to longer-latency responses. **C)** During hindlimb-only locomotion in the spinal state, SP afferents make synaptic contacts with lumbar circuits to evoke short-latency reflexes but ascending and descending pathways to and from the brain/cervical cord are abolished and no longer contribute to longer-latency responses.

### Concluding remarks

In conclusion, our results suggest that forelimb movements have a strong influence on hindlimb cutaneous reflexes and their modulation during quadrupedal locomotion. Additionally, some changes in reflex responses observed in spinal cats can be explained by the absence of forelimb movements, and not just because of removing supraspinal/cervical contributions. Humans are thought to have conserved a quadrupedal-like coordination of their arms and legs during walking (Pearcey and Zehr 2019) and making use of this interlimb circuitry could benefit rehabilitation approaches in various neurological conditions, such as stroke, Parkinson’s disease and spinal cord injury (Kaupp et al. 2018; Kawashima et al. 2008; Zhou et al. 2018). Animal models, such as the cat, provide a substrate to better understand the neural pathways and mechanical properties that help coordinate the limbs during locomotion and shape patterns of activity (Danner et al. 2017; Hurteau et al. 2018; Zhang et al. 2022).

## DATA AVAILABILITY STATEMENT

The datasets generated during and/or analysed during the current study are available from the corresponding author on reasonable request.

## ACKNOWLEDGEMENTS

We thank Philippe Drapeau (Université de Montréal) from the Rossignol and Drew labs for developing the data collection and analysis software.

## GRANTS

This work was supported by grants from the Natural Sciences and Engineering Research Council of Canada Grant NSERC RGPIN-2016-03790 (to A.F.) and the National Institutes of Health Grant R01 NS110550 (to A.F., I.A.R., and B.I.P.). A.F. and C.I-M. are Senior and Junior Research Scholars, respectively, supported by the Fonds de recherche du Québec – Santé (FRQS). J.H., J.A. and O.E. received doctoral scholarships from the FRQS. P.J. received a postdoctoral fellowship from the Centre de Recherche du CHUS.

## DISCLOSURES

No conflicts of interest, financial or otherwise, are declared by the authors.

## AUTHOR CONTRIBUTIONS

Author contributions: J.H. and A.F. designed research; J.H., R.A.A., S.M., S.Y., O.E., P.J., J.A., and C.L.. performed research; J.H., R.A.A., S.M., S.Y. and O.E. analyzed data; J.H., C.I.M., B.I.P., I.A.R. and A.F. wrote the paper.

## Notes

### Competing Interest Statement

The authors have declared no competing interest.

## REFERENCES

Abraham LD, Marks WB, Loeb GE. The distal hindlimb musculature of the cat. Cutaneous reflexes during locomotion. Exp Brain Res 58: 594–603, 1985.

Akay T, McVea DA, Tachibana A, Pearson KG. Coordination of fore and hind leg stepping in cats on a transversely-split treadmill. Exp Brain Res 175: 211–222, 2006.

Bernard G, Bouyer L, Provencher J, Rossignol S. Study of cutaneous reflex compensation during locomotion after nerve section in the cat. J Neurophysiol 97: 4173–4185, 2007.

Bretzner F, Drew T. Motor cortical modulation of cutaneous reflex responses in the hindlimb of the intact cat. J Neurophysiol 94: 673–687, 2005.

Buford JA, Smith JL. Adaptive control for backward quadrupedal walking. III. Stumbling corrective reactions and cutaneous reflex sensitivity. J Neurophysiol 70: 1102–1114, 1993.

Burke D. Clinical uses of H reflexes of upper and lower limb muscles. Clin Neurophysiol Pract 1: 9–17, 2016.

Capaday C, Stein RB. A method for simulating the reflex output of a motoneuron pool. J Neurosci Methods 21: 91–104, 1987.

Cruse H, Warnecke H. Coordination of the legs of a slow-walking cat. Exp Brain Res 89: 147–156, 1992.

Dambreville C, Labarre A, Thibaudier Y, Hurteau M-F, Frigon A. The spinal control of locomotion and step-to-step variability in left-right symmetry from slow to moderate speeds. J Neurophysiol 114: 1119–1128, 2015.

Danner SM, Shevtsova NA, Frigon A, Rybak IA. Computational modeling of spinal circuits controlling limb coordination and gaits in quadrupeds. Elife 6: e31050, 2017.

Delwaide PJ, Figiel C, Richelle C. Influence of the position of the upper limb on the excitability of the reflex arc of the soleus muscle. Electromyogr Clin Neurophysiol 13: 515–523, 1973.

Delwaide PJ, Figiel C, Richelle C. Effects of postural changes of the upper limb on reflex transmission in the lower limb. Cervicolumbar reflex interactions in man. J Neurol Neurosurg Psychiatry 40: 616–621, 1977.

Drew T, Rossignol S. A kinematic and electromyographic study of cutaneous reflexes evoked from the forelimb of unrestrained walking cats. J Neurophysiol 57: 1160–1184, 1987.

Duysens J, Loeb GE. Modulation of ipsi- and contralateral reflex responses in unrestrained walking cats. J Neurophysiol 44: 1024–1037, 1980.

Duysens J, Stein RB. Reflexes induced by nerve stimulation in walking cats with implanted cuff electrodes. Exp Brain Res 32: 213–224, 1978.

Eke-Okoro ST. Evidence of interaction between human lumbosacral and cervical neural networks during gait. Electromyogr Clin Neurophysiol 34: 345–349, 1994.

Ferris DP, Huang HJ, Kao P-C. Moving the arms to activate the legs. Exerc Sport Sci Rev 34: 113–120, 2006.

Forssberg H. Stumbling corrective reaction: a phase-dependent compensatory reaction during locomotion. J Neurophysiol 42: 936–953, 1979.

Forssberg H, Grillner S, Rossignol S. Phase dependent reflex reversal during walking in chronic spinal cats. Brain Res 85: 103–107, 1975.

Forssberg H, Grillner S, Rossignol S. Phasic gain control of reflexes from the dorsum of the paw during spinal locomotion. Brain Res 132: 121–139, 1977.

Frigon A, Akay T, Prilutsky BI. Control of Mammalian Locomotion by Somatosensory Feedback. Compr Physiol 12: 2877–2947, 2021.

Frigon A, Barrière G, Leblond H, Rossignol S. Asymmetric changes in cutaneous reflexes after a partial spinal lesion and retention following spinalization during locomotion in the cat. J Neurophysiol 102: 2667–2680, 2009.

Frigon A, Collins DF, Zehr EP. Effect of rhythmic arm movement on reflexes in the legs: modulation of soleus H-reflexes and somatosensory conditioning. J Neurophysiol 91: 1516–1523, 2004.

Frigon A, D’Angelo G, Thibaudier Y, Hurteau M-F, Telonio A, Kuczynski V, Dambreville C. Speed-dependent modulation of phase variations on a step-by-step basis and its impact on the consistency of interlimb coordination during quadrupedal locomotion in intact adult cats. J Neurophysiol 111: 1885–1902, 2014.

Frigon A, Rossignol S. Plasticity of reflexes from the foot during locomotion after denervating ankle extensors in intact cats. J Neurophysiol 98: 2122–2132, 2007.

Frigon A, Rossignol S. Short-latency crossed inhibitory responses in extensor muscles during locomotion in the cat. J Neurophysiol 99: 989–998, 2008a.

Frigon A, Rossignol S. Adaptive changes of the locomotor pattern and cutaneous reflexes during locomotion studied in the same cats before and after spinalization. J Physiol 586: 2927–2945, 2008b.

Haridas C, Zehr EP. Coordinated interlimb compensatory responses to electrical stimulation of cutaneous nerves in the hand and foot during walking. J Neurophysiol 90: 2850–2861, 2003.

Harnie J, Audet J, Klishko AN, Doelman A, Prilutsky BI, Frigon A. The Spinal Control of Backward Locomotion. J Neurosci 41: 630–647, 2021.

Harnie J, Audet J, Mari S, Lecomte CG, Merlet AN, Genois G, Rybak IA, Prilutsky BI, Frigon A. State- and Condition-Dependent Modulation of the Hindlimb Locomotor Pattern in Intact and Spinal Cats Across Speeds. Front Syst Neurosci 16: 814028, 2022.

Harnie J, Côté-Sarrazin C, Hurteau M-F, Desrochers E, Doelman A, Amhis N, Frigon A. The modulation of locomotor speed is maintained following partial denervation of ankle extensors in spinal cats. J Neurophysiol 120: 1274–1285, 2018.

Hiraoka K. Phase-dependent modulation of the soleus H-reflex during rhythmical arm swing in humans. Electromyogr Clin Neurophysiol 41: 43–47, 2001.

Hiraoka K, Nagata A. Modulation of the soleus H reflex with different velocities of passive movement of the arm. Electromyogr Clin Neurophysiol 39: 21–26, 1999.

Hundza SR, Zehr EP. Suppression of soleus H-reflex amplitude is graded with frequency of rhythmic arm cycling. Exp Brain Res 193: 297–306, 2009.

Hurteau M-F, Frigon A. A Spinal Mechanism Related to Left-Right Symmetry Reduces Cutaneous Reflex Modulation Independently of Speed During Split-Belt Locomotion. J Neurosci 38: 10314–10328, 2018.

Hurteau M-F, Thibaudier Y, Dambreville C, Chraibi A, Desrochers E, Telonio A, Frigon A. Nonlinear Modulation of Cutaneous Reflexes with Increasing Speed of Locomotion in Spinal Cats. J Neurosci 37: 3896– 3912, 2017.

Hurteau M-F, Thibaudier Y, Dambreville C, Danner SM, Rybak IA, Frigon A. Intralimb and Interlimb Cutaneous Reflexes during Locomotion in the Intact Cat. J Neurosci 38: 4104–4122, 2018.

Kagamihara Y, Hayashi A, Masakado Y, Kouno Y. Long-loop reflex from arm afferents to remote muscles in normal man. Exp Brain Res 151: 136–144, 2003.

Kao P-C, Ferris DP. The effect of movement frequency on interlimb coupling during recumbent stepping. Motor Control 9: 144–163, 2005.

Kaupp C, Pearcey GEP, Klarner T, Sun Y, Cullen H, Barss TS, Zehr EP. Rhythmic arm cycling training improves walking and neurophysiological integrity in chronic stroke: the arms can give legs a helping hand in rehabilitation. J Neurophysiol 119: 1095–1112, 2018.

Kawashima N, Nozaki D, Abe MO, Nakazawa K. Shaping appropriate locomotive motor output through interlimb neural pathway within spinal cord in humans. J Neurophysiol 99: 2946–2955, 2008.

Klishko AN, Akyildiz A, Mehta-Desai R, Prilutsky BI. Common and distinct muscle synergies during level and slope locomotion in the cat. J Neurophysiol 126: 493–515, 2021.

LaBella LA, Niechaj A, Rossignol S. Low-threshold, short-latency cutaneous reflexes during fictive locomotion in the “semi-chronic” spinal cat. Exp Brain Res 91: 236–248, 1992.

Lam T, Wolstenholme C, van der Linden M, Pang MYC, Yang JF. Stumbling corrective responses during treadmill-elicited stepping in human infants. J Physiol 553: 319–331, 2003.

Lecomte CG, Mari S, Audet J, Yassine S, Merlet AN, Morency C, Harnie J, Beaulieu C, Gendron L, Frigon A. Neuromechanical Strategies for Obstacle Negotiation during Overground Locomotion following Incomplete Spinal Cord Injury in Adult Cats. J Neurosci 43: 5623–5641, 2023.

de Leon RD, Hodgson JA, Roy RR, Edgerton VR. Locomotor capacity attributable to step training versus spontaneous recovery after spinalization in adult cats. J Neurophysiol 79: 1329–1340, 1998.

Loeb GE. The distal hindlimb musculature of the cat: interanimal variability of locomotor activity and cutaneous reflexes. Exp Brain Res 96: 125–140, 1993.

Mari S, Lecomte CG, Merlet AN, Audet J, Harnie J, Rybak IA, Prilutsky BI, Frigon A. A sensory signal related to left-right symmetry modulates intra- and interlimb cutaneous reflexes during locomotion in intact cats. Front Syst Neurosci 17: 1199079, 2023.

Matthews PB. Observations on the automatic compensation of reflex gain on varying the pre-existing level of motor discharge in man. J Physiol 374: 73–90, 1986.

Meinck HM, Piesiur-Strehlow B. Reflexes evoked in leg muscles from arm afferents: a propriospinal pathway in man? Exp Brain Res 43: 78–86, 1981.

Merlet AN, Jéhannin P, Mari S, Lecomte CG, Audet J, Harnie J, Rybak IA, Prilutsky BI, Frigon A. Sensory Perturbations from Hindlimb Cutaneous Afferents Generate Coordinated Functional Responses in All Four Limbs during Locomotion in Intact Cats. eNeuro 9: ENEURO.0178-22.2022, 2022.

Miller S, Ruit JB, Van der Meché FG. Reversal of sign of long spinal reflexes dependent on the phase of the step cycle in the high decerebrate cat. Brain Res 128: 447–459, 1977.

Pearcey GEP, Zehr EP. We Are Upright-Walking Cats: Human Limbs as Sensory Antennae During Locomotion. Physiology (Bethesda) 34: 354–364, 2019.

Percie du Sert N, Hurst V, Ahluwalia A, Alam S, Avey MT, Baker M, Browne WJ, Clark A, Cuthill IC, Dirnagl U, Emerson M, Garner P, Holgate ST, Howells DW, Karp NA, Lazic SE, Lidster K, MacCallum CJ, Macleod M, Pearl EJ, Petersen OH, Rawle F, Reynolds P, Rooney K, Sena ES, Silberberg SD, Steckler T, Würbel H. The ARRIVE guidelines 2.0: Updated guidelines for reporting animal research. PLoS Biol 18: e3000410, 2020.

Potocanac Z, Pijnappels M, Verschueren S, van Dieën J, Duysens J. Two-stage muscle activity responses in decisions about leg movement adjustments during trip recovery. J Neurophysiol 115: 143–156, 2016.

Pratt CA, Chanaud CM, Loeb GE. Functionally complex muscles of the cat hindlimb. IV. Intramuscular distribution of movement command signals and cutaneous reflexes in broad, bifunctional thigh muscles. Exp Brain Res 85: 281–299, 1991.

Prochazka A, Sontag KH, Wand P. Motor reactions to perturbations of gait: proprioceptive and somesthetic involvement. Neurosci Lett 7: 35–39, 1978.

Pruszynski JA, Scott SH. Optimal feedback control and the long-latency stretch response. Exp Brain Res 218: 341–359, 2012.

Quevedo J, Stecina K, Gosgnach S, McCrea DA. Stumbling corrective reaction during fictive locomotion in the cat. J Neurophysiol 94: 2045–2052, 2005a.

Quevedo J, Stecina K, McCrea DA. Intracellular analysis of reflex pathways underlying the stumbling corrective reaction during fictive locomotion in the cat. J Neurophysiol 94: 2053–2062, 2005b.

Sakamoto M, Endoh T, Nakajima T, Tazoe T, Shiozawa S, Komiyama T. Modulations of interlimb and intralimb cutaneous reflexes during simultaneous arm and leg cycling in humans. Clin Neurophysiol 117: 1301–1311, 2006.

Sakamoto M, Tazoe T, Nakajima T, Endoh T, Komiyama T. Leg automaticity is stronger than arm automaticity during simultaneous arm and leg cycling. Neurosci Lett 564: 62–66, 2014.

Sakamoto M, Tazoe T, Nakajima T, Endoh T, Shiozawa S, Komiyama T. Voluntary changes in leg cadence modulate arm cadence during simultaneous arm and leg cycling. Exp Brain Res 176: 188–192, 2007.

Schillings AM, Mulder T, Duysens J. Stumbling over obstacles in older adults compared to young adults. J Neurophysiol 94: 1158–1168, 2005.

Schillings AM, Van Wezel BM, Duysens J. Mechanically induced stumbling during human treadmill walking. J Neurosci Methods 67: 11–17, 1996.

Schillings AM, van Wezel BM, Mulder T, Duysens J. Muscular responses and movement strategies during stumbling over obstacles. J Neurophysiol 83: 2093–2102, 2000.

Shimamura M, Aoki M. Effects of spino-bulbo-spinal reflex volleys on flexor motoneurons of hindlimb in the cat. Brain Res 16: 333–349, 1969.

Shimamura M, Mori S, Yamauchi T. Effects of spino-bulbo-spinal reflex volleys on extensor motoneurons of hindlimb in cats. J Neurophysiol 30: 319–332, 1967.

Smith JL, Carlson-Kuhta P, Trank TV. Forms of forward quadrupedal locomotion. III. A comparison of posture, hindlimb kinematics, and motor patterns for downslope and level walking. J Neurophysiol 79: 1702–1716, 1998.

Thibaudier Y, Frigon A. Spatiotemporal control of interlimb coordination during transverse split-belt locomotion with 1:1 or 2:1 coupling patterns in intact adult cats. J Neurophysiol 112: 2006–2018, 2014.

Thibaudier Y, Hurteau M-F, Telonio A, Frigon A. Coordination between the fore- and hindlimbs is bidirectional, asymmetrically organized, and flexible during quadrupedal locomotion in the intact adult cat. Neuroscience 240: 13–26, 2013.

Van Wezel BM, Ottenhoff FA, Duysens J. Dynamic control of location-specific information in tactile cutaneous reflexes from the foot during human walking. J Neurosci 17: 3804–3814, 1997.

Wand P, Prochazka A, Sontag KH. Neuromuscular responses to gait perturbations in freely moving cats. Exp Brain Res 38: 109–114, 1980.

Zehr EP, Collins DF, Chua R. Human interlimb reflexes evoked by electrical stimulation of cutaneous nerves innervating the hand and foot. Exp Brain Res 140: 495–504, 2001.

Zehr EP, Duysens J. Regulation of arm and leg movement during human locomotion. Neuroscientist 10: 347– 361, 2004.

Zehr EP, Komiyama T, Stein RB. Cutaneous reflexes during human gait: electromyographic and kinematic responses to electrical stimulation. J Neurophysiol 77: 3311–3325, 1997.

Zehr EP, Stein RB. What functions do reflexes serve during human locomotion? Prog Neurobiol 58: 185–205, 1999.

Zhang H, Shevtsova NA, Deska-Gauthier D, Mackay C, Dougherty KJ, Danner SM, Zhang Y, Rybak IA. The role of V3 neurons in speed-dependent interlimb coordination during locomotion in mice. Elife 11: e73424, 2022.

Zhou R, Parhizi B, Assh J, Alvarado L, Ogilvie R, Chong SL, Mushahwar VK. Effect of cervicolumbar coupling on spinal reflexes during cycling after incomplete spinal cord injury. J Neurophysiol 120: 3172–3186, 2018.

